# A taxonomy for human social perception: Data-driven modelling with cinematic stimuli

**DOI:** 10.1101/2023.09.28.559888

**Authors:** Severi Santavirta, Tuulia Malén, Asli Erdemli, Lauri Nummenmaa

## Abstract

Every day, humans encounter complex social situations that need to be encoded effectively to allow interaction with others. Yet, principles for organizing the perception of social features from the external world remain poorly characterised. In this large-scale study we investigated the principles of social perception in dynamic scenes. In the primary dataset, we presented 234 movie clips (41 minutes) containing various social situations to 1140 participants and asked them to evaluate the presence of 138 social features in each clip. Analyses of the social feature ratings revealed that some features are perceived categorically (present or absent) and others continuously (intensity) and simple social features requiring immediate response are perceived most consistently across participants. To establish the low-dimensional perceptual organization for social features based on movies, we used principal coordinate analysis and consensus clustering for the feature ratings. These dimension reduction analyses revealed that the social perceptual structure can be modelled with eight main dimensions and that behaviourally relevant perceptual categories emerge from these main dimensions. This social perceptual structure generalized from the perception of unrelated Hollywood movie clips to the perception of a full Finnish movie (70 min), and to the perception of static images (n = 468) and across three independent sets of participants (n = 2254). Based on the results, we propose **eight basic dimensions of social perception** as a model for rapid social perception where social situations are perceived along eight orthogonal perceptual dimensions (most importantly emotional valence, empathy versus dominance, and cognitive versus physical behaviour).

## Introduction

Social interaction is central to humans. We live in complex social systems (Tomasello, 2014) and both cooperation and social competition require perception, interpretation, and prediction of other people’s actions, goals, and motives. Accordingly, social signals are granted priority in the attention circuits, and a large bulk of data show that human faces and bodies are rapidly detected and prioritized in natural scenes (Fletcher-Watson et al., 2008; Ro et al., 2007) underlining the importance of social information perception for humans. Dynamic interactive theory of person construal (DI) proposes that social perception is a dynamic interaction of low-level sensory information perception and high-level cognitive processes (Freeman & Ambady, 2011). Low-level sensory information, most importantly vision and hearing, is limited by the constraints of our senses while high-level cognitive processes in social perception involve, for example, our prior experiences, goals and motives, and the current emotional state. These two parallel processing streams influence each other bidirectionally and dynamically shaping the way we perceive the social world around us.

Despite the complexity, dimensionality and fast temporal scale of social processes (Adolphs et al., 2016), humans are capable of perceiving instantly multiple simultaneously occurring social features unfolding in distinct temporal time scales, ranging from other people’s identities, intentions and actions to the subtle affective characteristics of social interaction and hierarchy in social groups. Given the computational limits of the human brain and the requirement for fast and flexible responses in social interaction, it is unlikely that all available social information can be perceived and interpreted independently all the time (Freeman et al., 2012). It is thus likely that the brain initially parses the social world along some higher-order primary dimensions, but the actual organization of these social perceptual dimensions remains poorly characterised.

Humans use heuristics to make fast and accurate judgements from incomplete information without complex computations (Gigerenzer & Brighton, 2009). The social perceptual system could use similar heuristics for rapidly extracting the main social information of the situation based on the associations between the perceived social features. These associations could be non-deliberative assessments of social features (e.g. seeing a shouting person, indicating that someone is angry or in distress) that help the perceiver to make fast inferences with minimal processing effort. The DI theory describes this possible heuristic as a hierarchy from cues (e.g. angry face) through categorization (e.g. angry person) to stereotyping people (e.g. hostile person) (Freeman & Ambady, 2011). With this framework, it is possible to explain how people can make such rapid judgements in complex social situations by inferring that through evolutionary or learned processes humans automatically connect easily and rapidly extracted social cues with more complex inferences of other people. There is also some evidence, that conceptual trait knowledge may be learned through social perception (Stolier et al., 2020).

The DI theory provides a conceptualization for social perception, but it does not describe what kind of social information is extracted to promote fast social evaluations. Various theoretical and data driven approaches have identified candidate categories and dimensions of social perception. Early factor analytic studies on semantic spaces revealed that semantic judgements vary on mostly three dimensions, valence, potency and activity (Osgood & Suci, 1955), which are also central to dimensional models of emotions such as the circumplex model of affect (Russell et al., 1989). The stereotype content model and dual perspective model of agency and communion, in parallel, describe that people and groups are perceived in two dimensions: warmth/communion and competence/agency (Abele & Wojciszke, 2014; Fiske, 2018). The ABC of stereotypes is a data driven extension for the warmth/communion and competence/agency model and it organizes people and social groups based on their agency/socioeconomic success, and conservative–progressive beliefs while communion is not and independent factor but rather a combination of these two dimensions (Koch et al., 2016). The 3d mind model, in turn, focuses on mental state concepts. It states that the mind organizes mental states in three dimensions: valence, rationality and social impact (Thornton & Tamir, 2020). Additionally, taxonomies have been developed for personality characteristics and person perception (Goldberg, 1990; Lee & Ashton, 2004; McCrae & Costa, 1987; Simms, 2007), human values (Schwartz et al., 2012) or goals (Wilkowski et al., 2020) social situations (Parrigon et al., 2017; Rauthmann et al., 2014) and action understanding (Thornton & Tamir, 2022).

A large bulk of studies has focused on studying trait inferences from standardized images of faces (Sutherland & Young, 2022). Valence and dominance (also named as trustworthiness and dominance or warmth and competence) have been identified as the two basic evaluative dimensions of faces (Lin et al., 2021; Oosterhof & Todorov, 2008). When perceiving faces of children (signalling no threat) the trustworthiness & dominance are replaced with niceness & shyness implying that impressions have a functional basis (Collova et al., 2019). The core valence – dominance model has since been extended with youthful-attractiveness dimension (Sutherland et al., 2013; Vernon et al., 2014). Sex characteristics masculinity and femininity have been usually considered a single continuum where femininity has been associated with youthful-attractiveness (Sutherland et al., 2020) or femininity has been considered separate from youthfulness (Lin et al., 2021) or masculinity has been coupled with dominance (Oosterhof & Todorov, 2008; Sutherland et al., 2013). In addition, personality traits are also associated with specific facial features, but the Big Five personality trait perception cannot be fully described with the two-dimensional valence and dominance model (Walker & Vetter, 2016). However, this type of perceptual studies fail to capture all important social perceptual information since some personality trait dimensions are inferred primarily from the bodies or from the whole person instead of the faces alone (Hu & O’Toole, 2023) and because facial images only convey a “snapshot” of the constantly evolving interactive signals that faces convey in real life.

Although social perception involves dynamic encoding of the interaction between the situation, people and their actions, prior studies have mapped the perception of these domains separately and complex real life social interaction has been simplified to static stimuli such as images, written sentences, or lexical similarities of words. While the previously described taxonomies of social perception cover different aspects of social cognition, many of them are however interlinked and overlapping (Horstmann et al., 2021; Lin & Thornton, 2023; Wilkowski et al., 2020) and the inferred social trait structure is shown to be highly similar across social cognitive domains, including face impressions, familiar person knowledge and group stereotypes (Stolier et al., 2020). Further, a recent study showed that mental state and trait inferences are not independent of each other in naturalistic perception (Lin & Thornton, 2023). Nevertheless, a psychological taxonomy integrating person’s traits, states, behaviour, and situation characteristics for dynamic social perception is currently lacking.

Fully realistic stimuli for every day social situations that would be suitable for controlled experiments is extremely difficult to obtain. Hence, movies have offered powerful means for studying social perception dynamically even though they typically present amplified or stereotypical social situations. Regardless of the divergence from real life, movie scenes typically present dynamic, socially rich episodes with high affective intensity and frequency. Consequently social neuroscience has successfully adopted movies for studying neural bases of social and emotional processing (Adolphs et al., 2016; Lahnakoski et al., 2012; Saarimäki, 2021; Santavirta et al., 2023). Large-scale brain networks are found to encode a limited number of primary socially relevant dimensions from dynamic video input consistently across individuals with dimension specific brain response patterns (Santavirta et al., 2023). Due to neuroimaging constraints the behavioural results from that study should be considered as preliminary since the study focused on a limited number of the most consistently perceived social features with limited participant pool. Nevertheless, the results confirmed that rapid social perception can be reduced to a limited set of theoretical social dimensions that capture different qualities of social situations. The present study builds on these findings using more robust set of participants, stimuli, and social features with more comprehensive analytical techniques to establish a reduced model for rapid social perception.

In addition to the dimensionality of social perception, it remains unresolved whether social perceptual processes rely on identification of certain social features or are the perceived intensities also evaluated. In many areas of perception, people tend to automatically assign the percept into distinct categories, even though the transitions are continuous. For example, colours, speech sounds and facial expressions are commonly automatically perceived categorically (Fugate, 2013). It is thus likely that categorical perception would also be common across other domains of social perception. To make quick judgements of the social situation, only the presence of certain features may be important to notice, but for other social features also the intensity may be relevant. Thus, the perceptual system may process some social features categorically (e.g. someone is talking or not) or continuously (e.g. how trustworthy someone is).

Finally, interindividual variation in the perception of various social features remains poorly characterized. For example, colour perception (Emery & Webster, 2019), susceptibility for visual illusions (Carbon, 2014) and facial expression recognition accuracy (Calvo & Nummenmaa, 2016) all vary substantially across individuals. Then, face perception studies have identified that people evaluate facial trustworthiness, dominance and attractiveness relatively consistently (but see (Hehman et al., 2017)) from standardized images (Carré et al., 2009; Jones et al., 2004; Oosterhof & Todorov, 2008) but it is yet to be established whether people perceive observable and inferred social features with similar consistency across individuals when perceiving dynamic social scenes.

## Overview of the studies

The objectives of the present studies were 1) to establish a low-dimensional taxonomy for rapid social perception based on movie stimuli, 2) to investigate the between-participant consistencies in the perception of different social features and 3) characterise whether perception for specific social features is categorical or continuous. We defined social perception broadly as the perception of all information extracted rapidly from other people that is potentially useful when interacting with them avoiding the separation of the persons, their actions, and situations. The following hypotheses were tested:

**Hypothesis 1**: Social perceptual features are perceived with variable agreement between perceivers and either categorically (present or absent) or continuously.

**Hypothesis 2**: Social perception is organized along a limited set of perceptual dimensions.

**Hypothesis 3**: Dimensionality in social perception generalizes across perceivers and conditions (generalizations across stimuli and stimulus types).

We tested these hypotheses in a series of four studies. We collected a large-scale online survey data where subjects (N = 1140) viewed short unrelated Hollywood movie clips (N = 234) with varying social content. Subjects were asked to evaluate the prominence of a large set (N = 138) of different social features in the videos. This primary dataset was used in all four studies. We then analysed how people perceive and evaluate different social features (Hypothesis 1, Study 1), and whether the high-dimensional social perceptual space can be modelled with a limited set of primary evaluative dimensions (Hypothesis 2, Study 2). Next, we tested for the generalizability of the resulting solution with a different movie stimulus and participants (Hypothesis 3, Study 3). An independent set of participants (N = 5) watched a full non-Hollywood movie as short clips and evaluated the presence of social features. Finally, we tested the generalizability of identified social perceptual model across stimulus types from the perception of dynamic movie clips to static images (Hypothesis 3, Study 4). To this end, we ran a large-scale online survey where subjects (N = 1109) viewed movie frames (N=468) extracted from the primary movie clip database and rated them similarly as in Experiment 1.

The results indicated that between-participant agreement of perception varied between different social features, and that individual social features are perceived either categorically (present or absent) or continuously (intensity). Rapid social perception in the movie stimuli could be modelled with eight orthogonal main dimensions (Pleasant – Unpleasant, Empathetic – Dominant, Physical – Cognitive, Disengaged – Loyal, Introvert – Extravert, Playful – Sexual, Alone – Together and Feminine – Masculine). This model generalized across perceivers, across distinct movie stimuli and from perception of dynamic movies to the perception of movie frames. Overall, between-participant agreement in social perception was higher when participants evaluated movies versus static images. Altogether the results establish a low-dimensional model of social perceptual processing, where emotional valence, dominance versus empathy, and cognitive functioning versus physical behaviour are the most fundamental dimensions.

**Figure 1.**
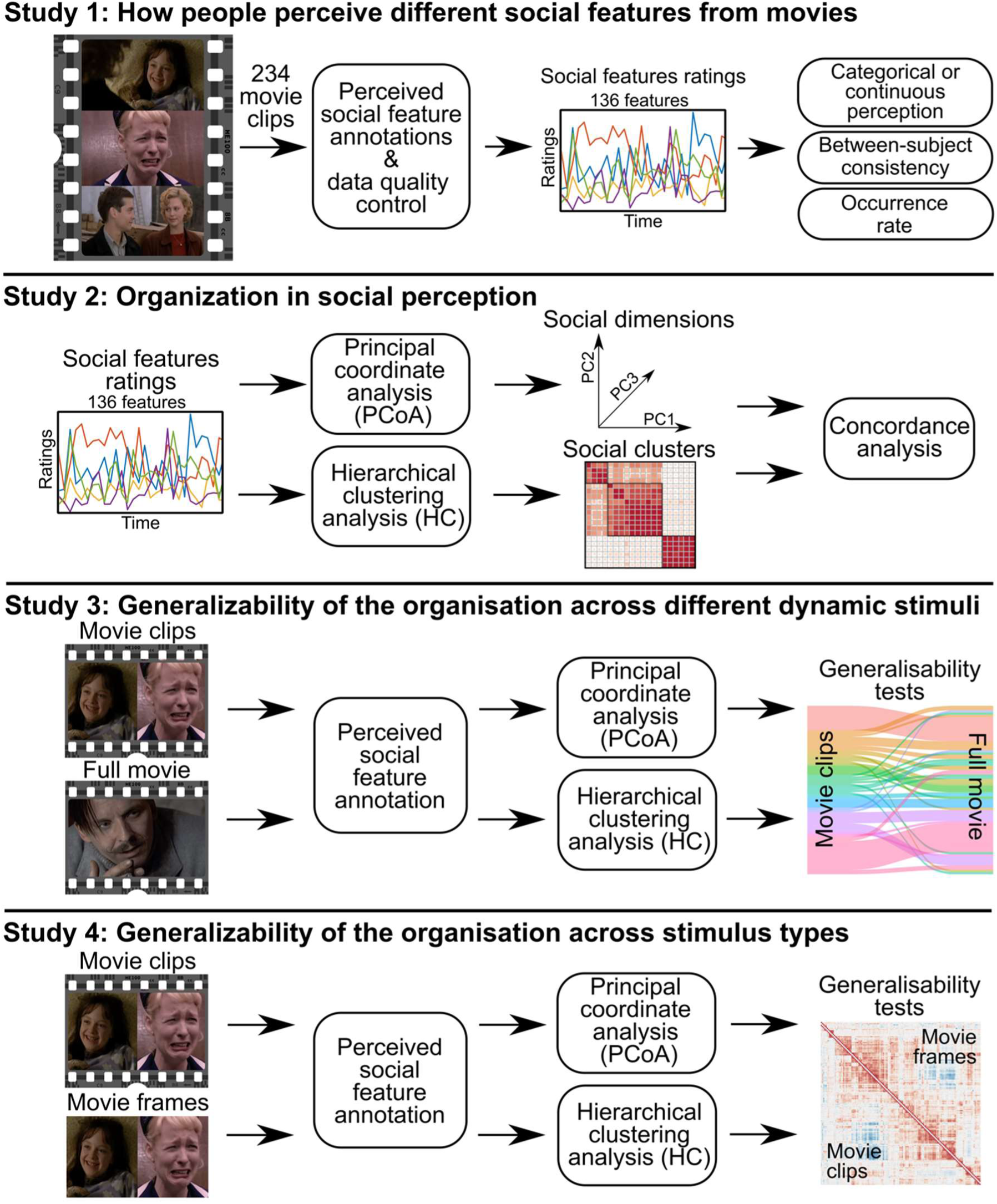
Overview of the studies. Study 1 investigated how people evaluate the presence of 136 social features in 234 movie clips. Study 2 mapped dimensional structure of social perception based on the ratings collected in Study 1. Study 3 tested the generalizability of the dimensional structure against independent dataset acquired while viewing a 70-minute feature film. Study 4 tested the generalisability of the organisation from the perception of dynamic videos to the perception of static images.

## Study 1

The purpose of the Study 1 was to identify how people evaluate the presence of different social features from dynamic movie stimulus. We hypothesized that some social features are primarily perceived as either present or absent (categorical perception) while for other features the intensity of the feature is perceived (continuous perception).

## Method

### Evaluated social features

Perceptual ratings were collected for a broad set of social features. We initially defined a set of social features for annotation that would be comprehensive but not overly exhaustive. Some studies have tackled the feature selection by letting participants spontaneously define the scales and dimensions (Koch et al., 2016; Nicolas et al., 2022; Osgood & Suci, 1955). Letting participants freely choose the rated features could however result in a bias towards conscious reasoning and overlook unconscious processing of important social features. Therefore, we used predefined perceptual features that described social perception across multiple levels.

The feature set was compiled based on existing theories of social perception. The linguistic category model proposes that interpersonal language contains verbs, and adjectives that describe persons and their interactions ranging from concreate observable actions to abstract inferred states (Semin & Fiedler, 1991). Interpersonal verbs may be further categorised into those describing states or behaviours of the sentence subject (such as kick, admire) or the states evoked in the sentence object (such as surprise, amaze) and the verbs may have positive or negative semantic valence (such as love, hate) (Semin & Fiedler, 1991). Hence, the broad categories should contain features described with adjectives and verbs in different levels of abstraction (observable actions – inferred states) and the broad categories should also cover the whole triad from the perception of person(s) to the perception situation and behaviour (Funder, 2006). Thus, we selected person’s traits, person’s physical characteristics, person’s internal situational states, somatic functions, sensory states, qualities of the social interaction, communicative signals and persons’ movement as the broad perceptual categories. Multiple social features should be selected from each of these categories.

Second, we searched the literature for related taxonomies for features that would describe a subset of the pre-defined broad social categories. In previous studies, social cognition and perception has been characterised across multiple levels. On the level of single individuals, these range from stereotype-like predispositions such as agency and communion or warmth and competence (Abele & Wojciszke, 2014; Fiske, 2018), through rapidly varying and observable psychological processes such as emotions (Russell et al., 1989) to relatively invariant characteristics of individuals such as personality traits (Goldberg, 1990; Lee & Ashton, 2004). Finally, some models characterise the situational psychological and social characteristics such as the person’s goals and values (Schwartz et al., 2012; Wilkowski et al., 2020) and the social situation the individual is embedded in (Parrigon et al., 2017; Rauthmann et al., 2014).

While these taxonomies describe regularities in the social domain, they describe specific areas such as personality, stereotypic evaluations of social groups, human values, and situations, none of them directly assesses the whole social perceptual domain from person perception the perception of social situations. In addition, the many of the described dimensions are highly correlated between different domains which indicates that affective-cognitive phenomena cannot be fully distinguished from each other (Horstmann et al., 2021; Wilkowski et al., 2020). Therefore, we did not test the generalizability of any single one of these taxonomies in the context of social perception, but instead we took an integrative approach by selecting evaluated social features from all these models. These taxonomies did not include features that would describe sensory or somatic functions of people or social communicative signals and hence we refined the evaluated set with features from these categories. The final list was based on an updated set of features used in a previous neuroimaging study of social perception (Santavirta et al., 2023) also guided by previous work in social perception from neuroscience perspective (Hudson et al., 2020; Tomi Karjalainen et al., 2017; Lahnakoski et al., 2012; Manninen et al., 2017; Nummenmaa et al., 2011, 2023; Putkinen et al., 2023). The final feature set included a total of 138 features which was considered broad enough to capture the most important perceptual variation in the social world. We however acknowledge that the feature set cannot likely describe *all* possible situations and features people may encounter in the social world.

The full list of tested features and their associations with existing cognitive-affective taxonomies is listed in **Table SI-1**. The list contained 31 features describing person’s traits (e.g. pleasant, agreeable, extravert, honest, intelligent, impulsive, and kind). We also included eight main physical characteristics (e.g. feminine, masculine, and old). Next, we selected 24 features describing person’s internal, situational states (e.g. feeling insecure, exerting self-control, and wanting something), eight features describing somatic functions (e.g. sweating, feeling ill, eating) and eight features from sensory states (e.g. feeling pain, listening to something) were also included. To broaden the scope to social interaction (rather than focusing on the individual agents), we included 31 features describing the qualities of the social interaction (e.g. hostile, sexual, playful, and equal) and 14 features describing social communicative signals (e.g. talking, nonverbal communication, laughing, hitting/hurting someone). Finally, we included 14 features describing the bodily and facial movements with high social relevance (e.g. moving their body, moving towards someone, jumping, making facial expressions).

### Stimulus set

The stimuli were short clips selected primarily from mainstream Hollywood movies (N=234). The average duration of the movie clips was 10.5 seconds (range: 4.1 – 27.9 sec) with a total duration of 41 minutes. The stimuli contained a previously validated set of socioemotional movies used in neuroimaging studies (T. Karjalainen et al., 2018; Tomi Karjalainen et al., 2017; Lahnakoski et al., 2012; Nummenmaa et al., 2021; Santavirta et al., 2023) and was complemented with similar movie clips selected from YouTube (https://www.youtube.com/) by three independent research assistants. The assistants were instructed to search for movie scenes with social content and cut them to approximately 10-second-long continuous scenes. The clips should present scenes spoken in English that could depict real life social interaction (science fiction or animation movies were excluded). The assistants were instructed to search for clips that would contain as variable social contents as possible. No further instructions of what social contents the scenes should contain were given to make sure that the principal researchers’ assumptions of the organization of social perception do not bias the stimulus selection. The clips depicted people from both sexes, from different age groups and multiple ethnicities. 118 clips contained females, 197 males, 77 children / adolescents, 192 adults, 10 older adults, 213 Caucasians, 44 African Americans, 10 persons from East Asia and 4 persons form the Middle East. See **Table SI-2** for short descriptions of each movie clip. This stimulus set and the acquired ratings were subsequently used in all the studies 1-4.

### Data collection

To ensure subject engagement and to reduce the cognitive demand of the study, each participant was randomized to evaluate a subset (39 clips) of the movie clips for a subset (6-8 features) of the social features (see section Experiment subsets in Supplementary materials). Participants were instructed to rate the perceived features in the video clip rather than their own internal states (such as emotions evoked by the films). They were asked to rate how much they perceived the given social feature from the movie clip by moving a slider on a continuous and abstract scale (VAS) from “absent” to “a lot”. The data were collected using online experiment platform Gorilla (https://gorilla.sc/) and the median experiment duration for the participants was 21 minutes and 18 seconds.

### Participants

Participants were recruited through Prolific (https://www.prolific.co/). Only fluent English-speaking adults using a computer or a tablet were allowed to participate. Concerns have been raised about the data quality in online studies (Webb & Tangney, 2022) and different recruiting platforms (Peer et al., 2022). Accordingly, we used strict data quality screening and participant inclusion criteria (see section Data quality control in Supplementary materials). New participants were recruited until 10 reliable ratings were obtained for each social feature per video clip (see section Sample size estimation in Supplementary materials). Altogether 1140 participants completed the study, and 44 (3,9 %) participants were excluded based on the quality control. The final dataset included 1 096 participants from 60 nationalities with over 327 000 data points. 515 participants were female (46,9 %) and the median age of the participants was 28 years (range: 18 – 78 years). The reported ethnicities were: White (788, 71.9 %), Black (136, 12.4 %), Mixed (81, 7.4%), Asian (47, 4.3 %) and Other (30, 2.7 %). See **Table SI-3** for the nationalities of participants.

### Analysing the distribution and consistency of social feature perception

Some of the rated social features are omnipresent in everyday social interaction (e.g. facial expressions) while others (e.g. hitting or hurting someone) occur infrequently. To investigate how often specific social features are present and whether the movie clips contained sufficient occurrences of the targeted social features, we calculated the feature occurrence rate. A feature was considered present in each movie clip if the average rating over annotators minus standard error (SEM) was above five (in the scale from 0 to 100) and occurrence rate was calculated as the proportion of all movie clips where the feature was present.

Some social features are inherently discrete and others continuous. For example, a person could be perceived as standing or not (categorical perception) whereas someone could be perceived as more or less trustworthy (continuous perception). To reveal how people evaluate the category and intensity of the social features, we analysed the shape of the population average distributions for the feature ratings. Bimodality coefficient (BC) and Hartigan’s dip test were used to test whether on average the people rate social features as unimodal or bimodal: Bimodality would suggest that the social feature is perceived categorically as either present or absent while unimodal distribution would indicate continuous intensity levels in perception. Hartigan’s dip test rejects unimodality at p < 0.05 and bimodality coefficient favours bimodality when BC > 0.555. Hartigan’s dip test is highly specific and less sensitive in identifying bimodality while BC is more sensitive but can also yield in false positive findings (Freeman & Dale, 2013). Hence, visual judgement of the distributions and the results of statistical tests should be considered complementary when assessing bimodality.

Finally, some social features are likely evaluated more consistently across observers than others. For example, people recognize others’ sex from their facial appearance with almost 100% accuracy (Bruce et al., 1993), whereas evaluation of personality-related features such as shyness is likely less consistent, particularly from the short stimulus clips. To assess the between-subject reliability in the ratings for each social feature we calculated intra-class correlation coefficients (ICC) (McGraw & Wong, 1996) for the feature ratings. Between-subject reliability was estimated by calculating the single measures ICC which measures the reliability of any single participants ratings compared to the others. A two-way random model with absolute agreement (ICC2) was selected for ICC calculations since both participants and movie clips are assumed to be randomly drawn from the underlying population of people and videos (Koo & Li, 2016). As previous studies on social perception have primarily relied on Cronbach’s alpha as consistency measure (Tavakol & Dennick, 2011), we also computed alphas for the sake of comparison.

## Results

### Distributions of social feature ratings

Figure 2 shows the distributions of population average ratings for all studied social features on the original VAS scale. Features “Coughing/sneezing” and “Vomiting/urinating/defecating” were excluded from the analyses because they were perceived present in less than 5% of the movie clips. Based on both the bimodality coefficient (highly sensitive for bimodality) and the Hartigan’s dip test (highly specific for bimodality) (**Figure SI-1**), the following features were considered as bimodal: talking, sitting, standing, making gaze contact, alone, feeling pain, feeling calm, and feminine. Additionally, bimodal coefficient favoured bimodality for other features also suggested by the visual representation of the features in Figure 2 (e.g. masculine, laying down, jumping, nude, sexual, kissing/hugging/cuddling, romantic, flirtative, touching someone, joking, laughing, eating/drinking, hungry/thirsty, tasting something, crying, yelling, hitting/hurting someone and physically aggressive, verbally aggressive).

**Figure 2.**
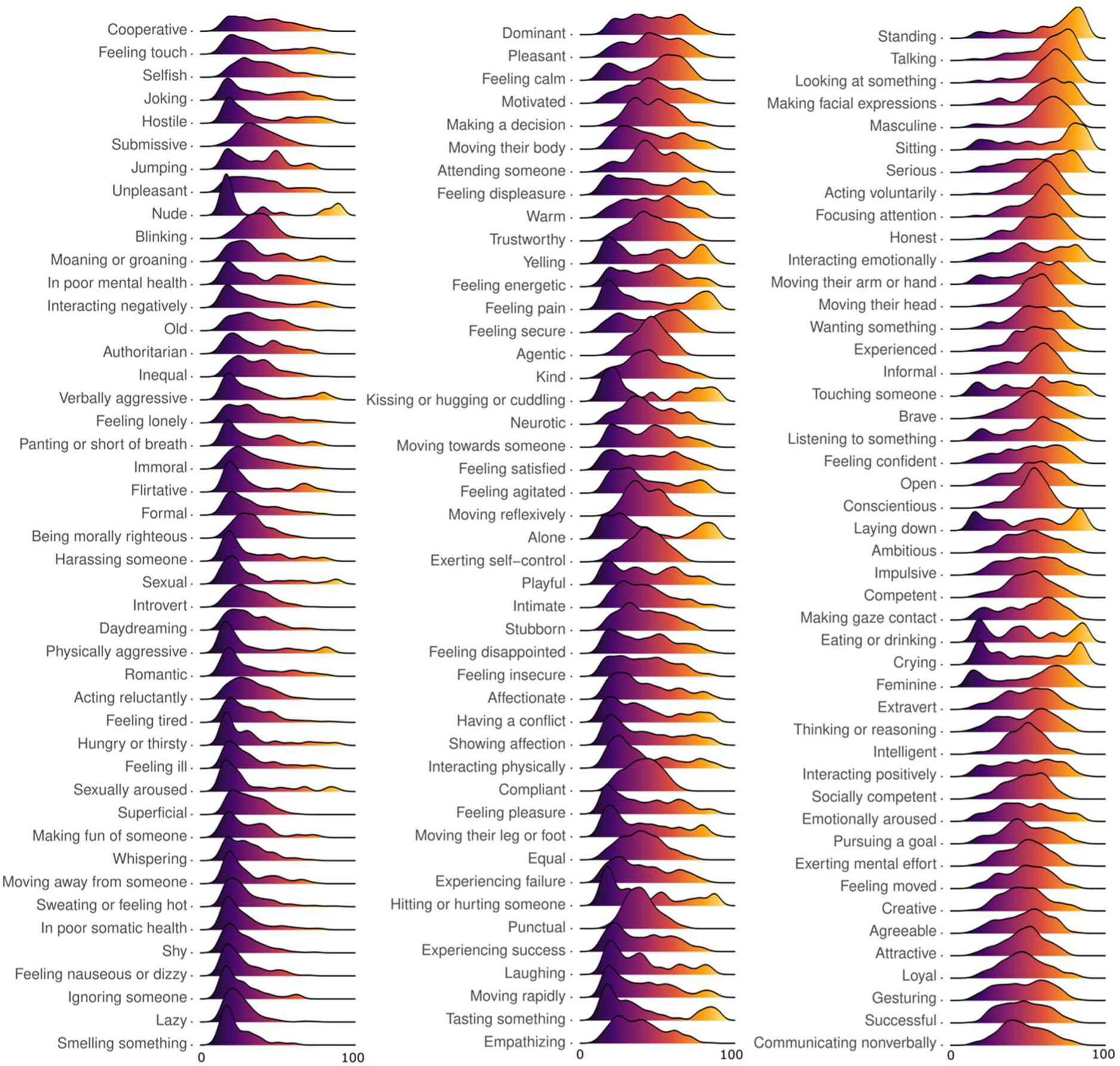
Distributions of the social feature ratings. The density plots visualise the estimated population level rating distributions for each feature (the rating from 0-100 in x-axis). Ratings under 5 (feature was not present) are not plotted for representative visualisation. The features are ordered based on the average intensity of the population level ratings.

### Between-subject reliability and occurrence rate of social feature ratings

Figure 3 visualises the feature-specific between-subject ICC:s, occurrence rates and bimodality scores. ICC:s for the ratings varied considerably between different social features (range: 0.08 – 0.75). Features with higher reliability (ICC > 0.6) related to concrete actions and interaction signals (e.g. sitting, eating, yelling, laughing, crying) and they were also typically perceived categorically (present or absent; Figure 3). Features with lower reliability (ICC < 0.2) mostly described subtle person-related characteristics (e.g. introvert, conscientious, compliant, exerting self-control). Overall ICC:s showed a decreasing gradient from actions perceived in categorical manner (e.g. standing, hitting or hurting someone), physical characteristics (e.g. feminine, masculine) and emotions (e.g. feeling pleasure, feeling satisfied, feeling pain) towards subtle characteristics related to personality and qualities of the social interaction (e.g. lazy, shy, superficial, informal, equal). Cronbach’s alphas ranged from 0.57 (punctual) to 0.97 (crying) and for 126 out of 136 features the Cronbach’s alpha was above 0.7.

**Figure 3.**
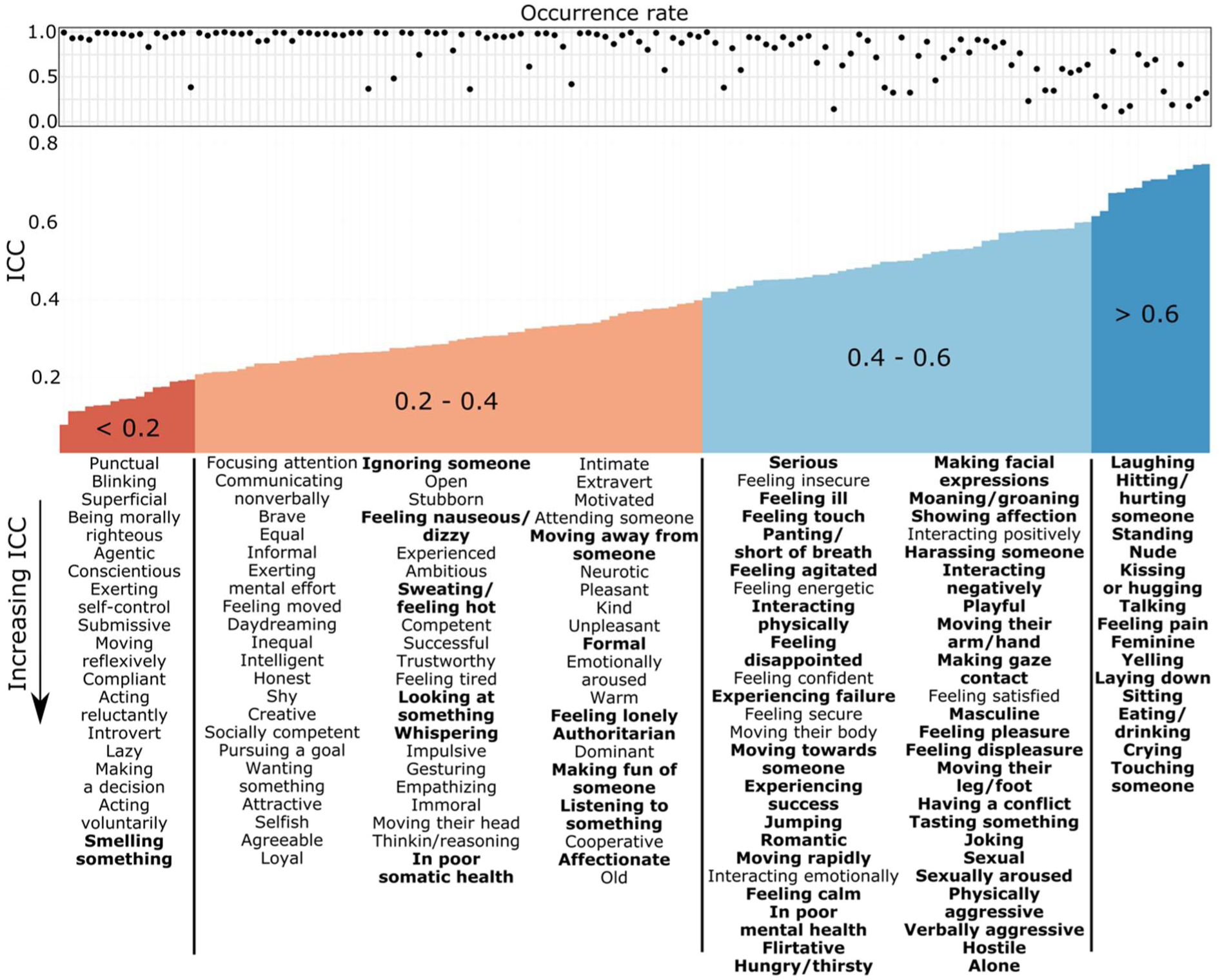
Feature occurrence rate, between-subject reliability, and bimodality. Pointplot visualises the occurrence rates (proportion of the movie clips) and barplot shows the distribution of the ICC:s for the feature ratings. The features below the plot are listed from the lowest (Punctual) to the highest (Touching someone) ICC. Categorically perceived social features are shown in boldface (BC > 0.555).

The mean occurrence rate for the analysed 136 features was 79% (range: 12% – 100%, Figure 3). Features with the lowest occurrence rate (< 20%) were binary features: nude, jumping, hitting or hurting someone, kissing or hugging, eating or drinking and laying down. While the occurrence rate was high for most of the social features, the rating intensities still varied between the movie clips indicating that stimuli presented variable social content to the participants (**Figure SI-2**).

### Interdependence between perceived frequency, between-subject reliability, and bimodality of the social features

Occurrence rates and ICC:s correlated negatively (r = –0.63, p < 0.001), occurrence rates and bimodality coefficients correlated negatively (r = –0.89, p < 0.001) and ICC:s correlated positively with bimodality coefficients (r = 0.84, p < 0.001) indicating strong associations between the measures. Less frequently present features were perceived with higher between-subject reliability than frequently present features. Features where bimodality tests favoured categorical perception were consistently perceived either absent or present while the continuously perceived features were less frequently completely absent. Finally, categorically perceived were evaluated with higher between-subject reliability than the continuously perceived features. Feature specific results for all outcome measures can be found from **Table SI-4.**

## Study 2

The purpose of Study 2 was to establish a low-dimensional taxonomy for social perception based on dynamic movie stimulus (Hypothesis 2). The dataset was the same that was used in Study 1. We used dimension reduction methods to filter the main information that the participant perceived from the total set of the 136 evaluated and present social features. Two different dimension reduction methods were selected to minimize the influence of the method to the results and to also get a more detailed description of the taxonomy.

## Method

### Stimuli, participants and evaluated social features

We used the full rating dataset from Study 1 thus the stimuli, participants and evaluated social features were the same as described in Study 1 methods.

### Low-dimensional model for social perception

We originally obtained ratings for the occurrence of a total of 136 social features. Because the stimulus space is high-dimensional, it is unlikely that all individual features would carry unique information independently of other simultaneously occurring features. We thus conducted two different dimension reduction analyses to build a low-dimensional model for social perception. Prior to the analyses, the ratings were averaged over subjects to estimate the population-level means. Standard errors (SEM) were used as a measure of the reliability of the averages. In the original rating scale (from 0 to 100) the average SEMs ranged from 2.4 (Nude) to 9.7 (Communicating nonverbally) which was considered as acceptable margin of error. A principal coordinate analysis (PCoA) (Gower, 1966) was used as a primary dimension reduction technique to establish the low-dimensional space of social perception in the movie stimulus. PCoA studies the dimensionality of data based on the covariance structure between the features. The main difference between principal coordinate analysis and principal component analysis (PCA) is the choice of distance metric while both produce orthogonal components that characterize the dimensionality of the underlying data structure. Our data contained social features that could be perceived either present or absent while some features may be perceived in every scene but with varying intensity. The Euclidean distance used by PCA would be insufficient in capturing the co-occurrence of these distinct data types, which motivated us to choose to study the covariance between the features with PCoA instead of the Euclidean distances.

PCoA can reveal orthogonal dimensions from any data, but the interpretation of the dimensions can be complicated because the components are mathematical and do not follow physiological or psychological principles underlying the original data. In addition, if people make fast judgements of situations relying on evaluations of just few basic dimensions, then more detailed social perceptual information should be identifiable by combining situational information from different orthogonal dimensions. To investigate this, we conducted a complementary consensus clustering analysis using hierarchical clustering (HC) (Murtagh & Contreras, 2012) to group social features into semantically distinct clusters that are not strictly orthogonal and finally investigated whether HC clusters can be constructed as combinations of the PCoA components.

#### Principal coordinate analysis

PCoA was implemented using R function cmdscale (https://www.rdocumentation.org/packages/stats/versions/3.6.2/topics/cmdscale) with a Pearson correlation matrix of individual social features as input. The statistical deviation from chance of each principal component’s (PC) eigenvalue and the loadings for social features for each component were tested with a permutation test. Component specific eigenvalue’s statistical deviation from chance would indicate that the component explains variation that is not found in random data further indicating that the dimension encodes meaningful variance.

Similarly, statistical deviation from zero in the loading of a social feature would indicate that the feature is significantly associated with the identified component. For each permutation round, a new dataset was generated by randomly shuffling the ratings of each social feature (columnwise) thereby breaking the synchrony between the columns of the data. Then a PCoA analysis was conducted for the shuffled dataset. The procedure was repeated 1 000 000 times to estimate the null distributions for the eigenvalues and the loadings of each PC. Finally, true eigenvalues of the PCs and the loadings of each PC were ranked to their corresponding null distributions to assess their statistical significance (exact p-value).

#### Consensus clustering analysis

Average hierarchical clustering (UPGMA) (Murtagh & Contreras, 2012) was used due to its simplicity and effectiveness in grouping social features (Santavirta et al., 2023). To establish stable and generalizable clustering results we used a consensus clustering approach from diceR package (Chiu & Talhouk, 2018) in R software (https://www.r-project.org/). Since we did not know how many stable clusters the evaluated social feature space would form, a consensus clustering result was obtained by subsampling 80% of the items from the total sample and then performing clustering in each subsample for different numbers of clusters (from 5 to 40). The subsampling was repeated 1000 times for each number of clusters to establish a consensus matrix over different numbers of clusters and over all subsamples. Finally, a representative and stable clustering result was obtained from the hierarchically ordered consensus matrix.

#### Comparison of PCoA and HC

Both PCoA and HC analyses reduce the social perceptual space based on the covariance structure between the rated social features and the results were expected to converge. To investigate how well the orthogonal components from PCoA link with the HC clusters we projected the candidate social features into 2D space using t-distributed stochastic neighbour embedding (t-SNE) which is a non-linear dimension reduction technique to visualise high-dimensional data in 2D surface (Van der Maaten & Hinton, 2008). The featurewise PC loadings for significant PCs were given as input for the t-SNE and hence each candidate social feature was mapped into 2D projection based on its loadings for significant PCoA components, rather than the raw rating values. If social features form separable groups similar with the HC clusters in the t-SNE projection it would indicate that the PCoA dimensions and HC clusters contain structural similarities and that the HC clusters can be described with the information from the PCoA dimensions. To assess the fine-grained connections between PCs and HC clusters, we obtained the mean PC loading for each cluster by averaging the feature PC loadings over all candidate features within the clusters. The statistical significance of the cluster loadings was then assessed similarly as the significance of the individual social features from the permuted null distributions. With this approach we could study in detail how each HC cluster could be described as combined information from the PCoA components.

#### Naming the identified social dimensions and clusters

We initially developed our own descriptors for the identified components and clusters, but to increase generalizability of the naming scheme, we also gathered candidate descriptors from researchers (N=10) and from the general population (N=92). Local participants included psychology/neuroscience researchers not involved in this study and the subjects for the general population sample were recruited through Prolific (https://www.prolific.com/). Only fluent English speakers from core English speaking countries (UK, USA, Canada, Australia and New Zealand) were recruited to ensure high language level of the participants. The components in Figure 4 and the lists or features within each cluster (Figure 5) were shown to participants one-by-one and the participants were asked to provide short primary and secondary descriptors for each of them. Researchers annotated both components and clusters, but a single online participant was randomized to name only components or clusters. The experiment was conducted in Gorilla (https://app.gorilla.sc/). Different forms of the same descriptive name were combined, and the frequencies of the unique descriptors were then calculated separately for the researchers and online participants. We also expected that large language models could perform well in describing the semantic similarities of different words. Hence, we also asked ChatGPT3.5 (https://chat.openai.com/) for the names and for short 1-2 sentence descriptions (See supplementary chatgtp_dimension_naming_conversation.zip for the exact prompts and responses). All the naming data were collected to **Table-SI5** (components) and **Table-SI6** (clusters) and the final names were defined as the consensus over the responses of authors, researchers, online participants and ChatGPT.

**Figure 4.**
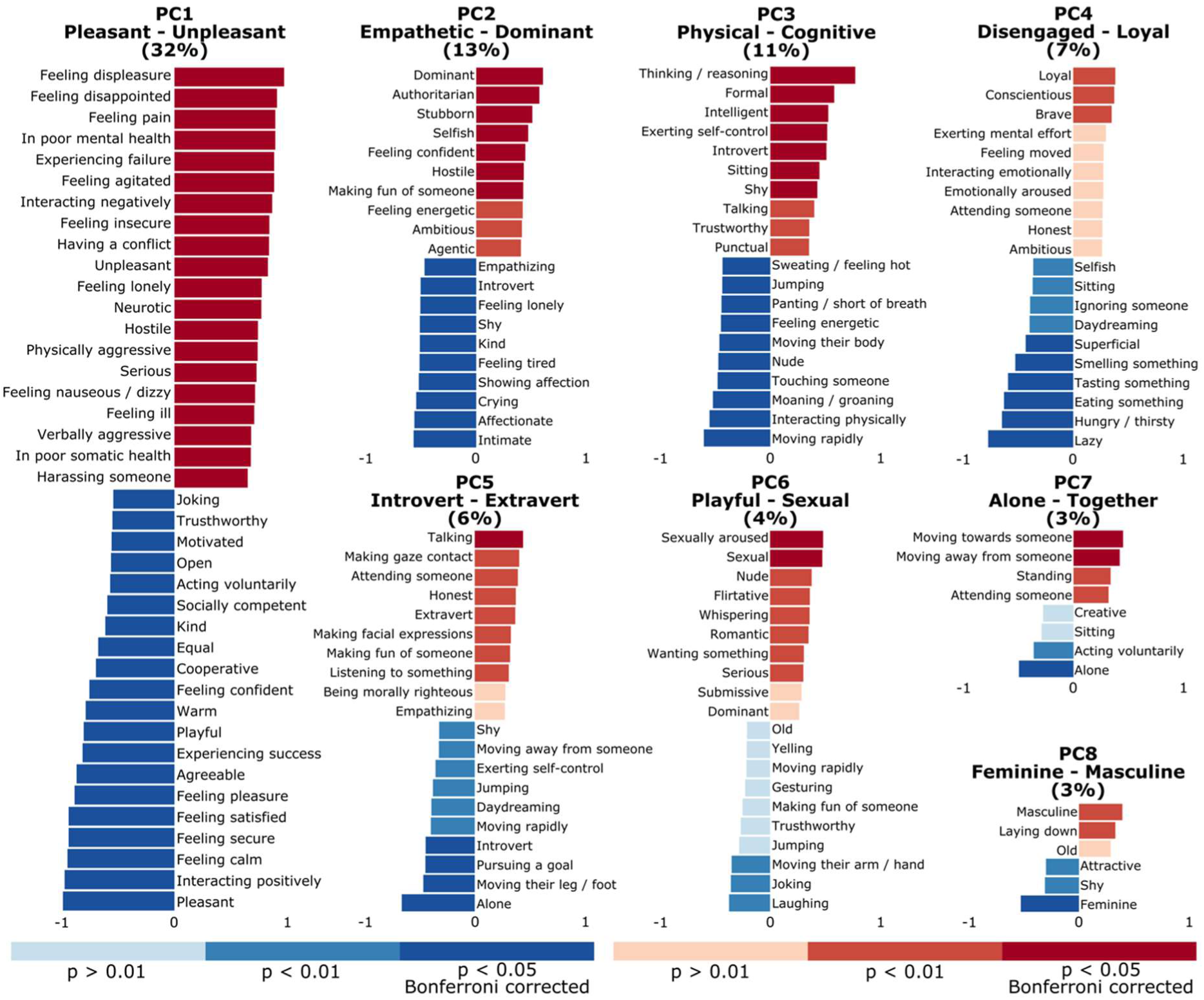
Results from the principal coordinate analysis. First eight principal components were statistically significant (p < 10^−7^). Barplots visualise the strongest loadings (correlations between the PC and the original features) for social features for each PCs. The names for the PCs describe the underlying perceptual social dimensions. Explained variance for each PC is shown in parentheses and the statistical significance thresholds for feature loadings are displayed with colour gradient.

**Figure 5.**
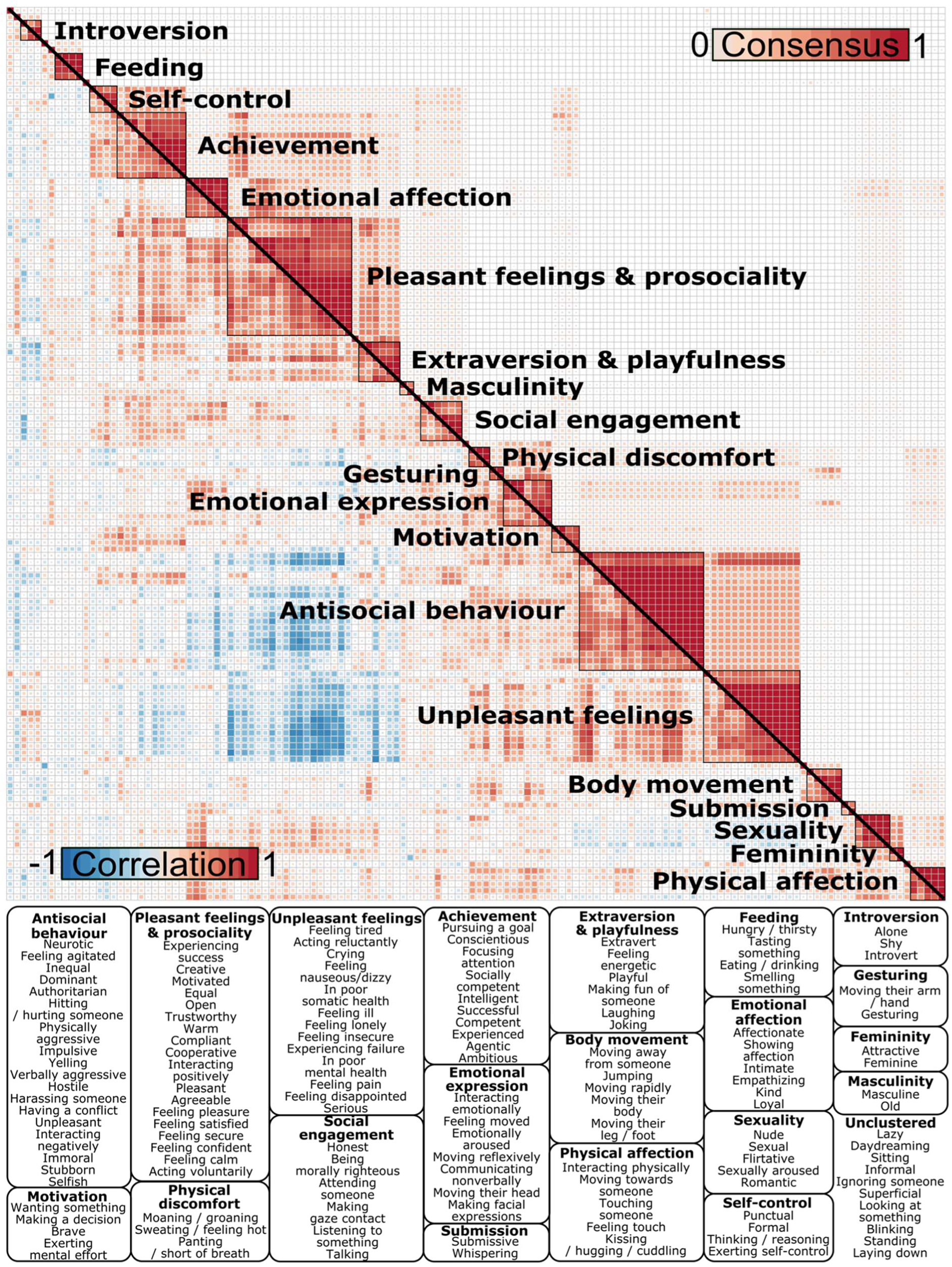
Results from the hierarchical clustering analysis. The lower triangle shows the correlation matrix of social features while the upper triangle shows the consensus matrix of the features from the consensus clustering analysis. A consensus matrix visualises how many times (out of all subsamples) each pair of features were clustered together. The boxes at the bottom show which individual social features were included in the identified clusters.

## Results

### Dimensional structure of social perception based the movie stimuli

Principal coordinate analysis revealed that social perceptual space of the movie stimuli can be reduced to eight statistically significant orthogonal principal components (p < 10^−7^, Figure 4). First three principal components (PCs) explained 55% and all eight PCs 78% of the total variance. PC1 (Pleasant – Unpleasant) ordered features based on their emotional valence from pleasant features “Pleasant”, “Interacting positively” and “Feeling calm” to unpleasant ones “Feeling displeasure”, “Feeling disappointed” and “Feeling pain”. PC2 (Empathetic – Dominant) related to the perceived dominance structure of the social interaction from dominant features “Dominance”, “Authority” and “Stubborn” to empathetic characteristics “Intimate”, “Affectionate” and “Crying”. PC3 (Physical – Cognitive) ordered features from physical engagement, energy consumption and impulsive behaviour (e.g. “Interacting physically”, “Nude”, “Feeling energetic”, “Moaning”, “Jumping”) to cognitive engagement and controlled behaviour (e.g. “Thinking or reasoning”, “Formal” and “Intelligent”). PC4 (Disengaged – Loyal) organized social perception from inactive self-related behaviour where people are disengaged from others (e.g. “Lazy”, “Superficial”, “Daydreaming”, “Eating something”, “Hungry / thirsty”) to engaged and proactive behaviours (e.g. “Conscientious”, “Loyal” and “Brave”). PC5 (Introvert – Extravert) related to the distinct social engagement and personality types, introversion (e.g. “Alone”, “Pursuing a goal” and “Introvert”) and extraversion (e.g. “Talking”, “Making gaze contact” and “Extravert”). PC6 (Playful – Sexual) described social contacts regarding their affiliative (e.g. “Joking” and “Laughing”) versus sexual nature (e.g. “Sexual”, “Sexually aroused” and “Nude”). PC7 (Alone – Together) described whether people were interacting with others or not and PC8 (Feminine – Masculine) described feminine and masculine characteristics. Beyond eight PCs, no component explained more variance than would be expected by chance even with a lenient thresholding (p < 0.05).

### Organization of social perception based on the consensus clustering analysis

Results for the consensus clustering (hierarchical algorithm; HC) are shown in Figure 5. The results confirmed that social perception is organized around the perceived emotional valence (negative correlation between pleasant and unpleasant features). However, consensus clustering further grouped emotionally valenced features into subclusters. Pleasant features formed three clusters (Pleasant feelings & prosociality, Emotional affection, and Extraversion & playfulness) and unpleasant features two clusters (Unpleasant feelings and Antisocial behaviour). Introversion formed one cluster and together with the cluster Extraversion & playfulness they established introversion – extraversion as a similar core social dimension as in the principal coordinate analysis. Feeding and Sexuality formed separate clusters, and communication-related features were clustered together into several clusters that described different communication types (Social engagement, Emotional expression, Gesturing and Physical affection). Clusters Self-control, Motivation and Achievement described traits of self-regulation, internal motivation and competence / success. Masculine and feminine characteristics formed independent clusters and additional clusters were identified for Body movement, Physical discomfort and Submission.

### Concordance between the PCoA and HC solutions

PCoA and HC revealed generally similar organization for social perceptual features, but the results were not identical due to different analytical approaches. For example, HC found semantically distinct clusters within emotionally valenced features while PCoA revealed the main dimension from unpleasant to pleasant features. The t-SNE suggested that the two alternative analytic solutions yield structural relationship since HC clusters formed clearly separable groups in this t-SNE space (Figure 6). HC clusters would have been unidentifiable in the t-SNE of PCoA feature loadings if there was no structural relationship between the solutions from the two methods.

**Figure 6.**
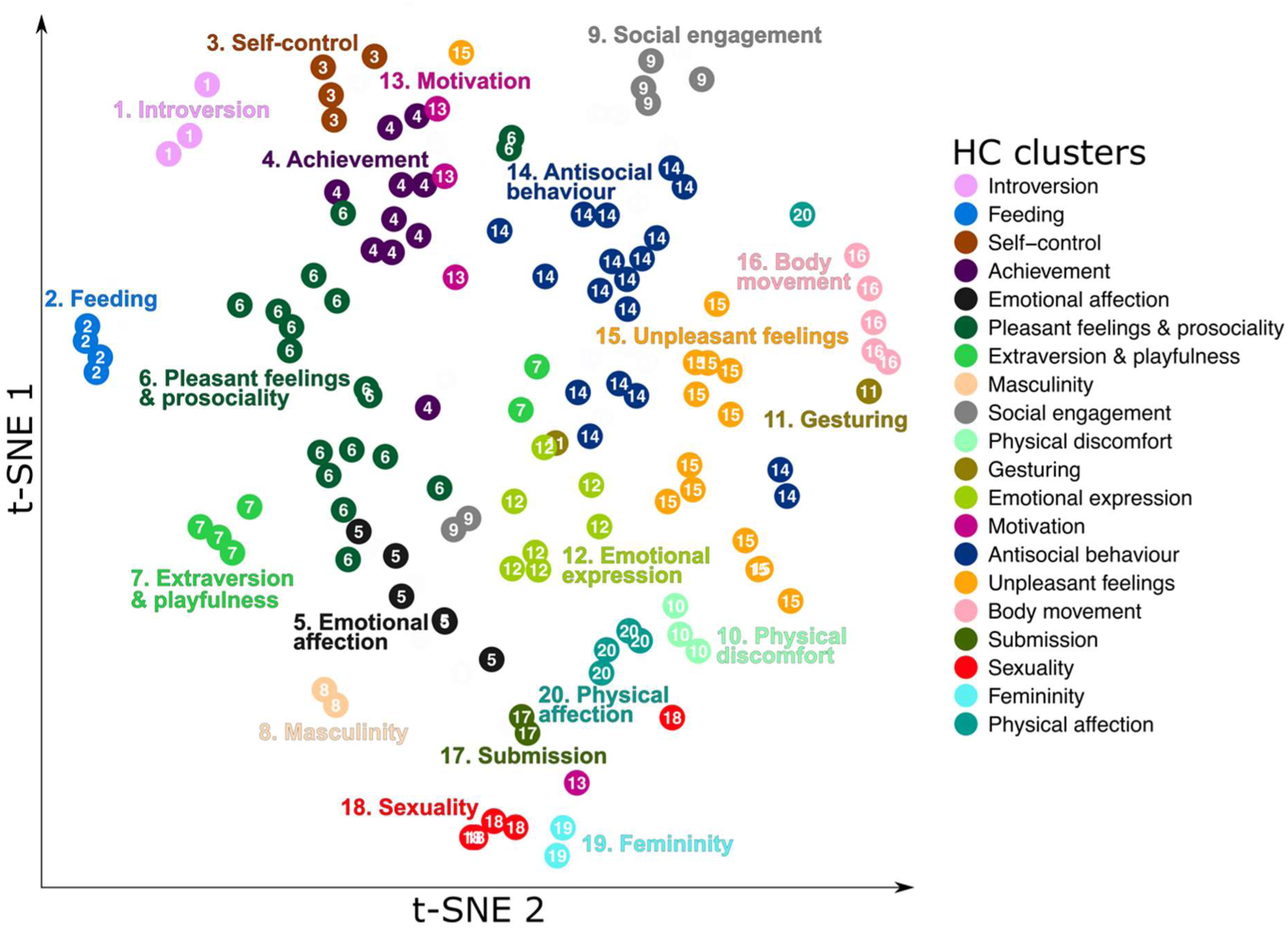
Relationship between HC and PCoA organizations for social perception. The t-SNE projection visualises candidate social features in 2D space based on their PCoA loadings for the eight significant main dimensions from PCoA. The colour and number of the social feature (point in the 2D space) mark which cluster the social feature was assigned to in the HC analysis and the cluster labels are visualised alongside the clusters. The visualisation shows that the HC clusters were well separated from each other based on the social feature loadings of the main dimensions in PCoA.

To further investigate the relationship between individual HC clusters and PCoA dimensions we obtained average PC loadings for each HC cluster (**Figure SI-3**). Some HC clusters loaded mainly on one corresponding main dimension in the PCoA solution, indicating that these perceptual dimensions were similarly captured by both analysis methods. The cluster Pleasant feelings & prosociality was based on features that loaded as pleasant in PC1 (Pleasant – Unpleasant), Unpleasant feelings from unpleasant features in PC1, Self-control from cognitive features in PC3 (Physical – Cognitive) and Feeding from the eating related features that loaded as inactive behaviours in PC4 (Disengaged – Loyal). Clusters Femininity and Masculinity loaded significantly on the opposite ends of PC8 (Feminine – Masculine). Overall, these results support that pleasantness, cognitive functioning, feeding and femininity-masculinity are perceptual main dimensions captured similarly by the alternative analysis methods.

HC did not find an independent cluster for dominant characteristics even though dominance was considered the second main dimension in PCoA. However, the concordance analysis revealed that pleasant and unpleasant characteristics formed semantically distinct clusters based on the perceived dominance structure supporting the existence of dominance as a perceptual dimension. Features in the cluster Antisocial behaviour loaded as unpleasant (PC1) and dominant (PC2) distinguishing antisocial behaviours from unpleasant feelings (unpleasant but not dominant characteristics). Similarly, features in the cluster Emotional affection loaded as pleasant (PC1) and empathic (PC2) while features in the cluster Extraversion & playfulness loaded as pleasant (PC1) and dominant (PC2) distinguishing these pleasantly perceived characteristics from the solely pleasant features in the cluster Pleasant feelings & prosociality (**Figure SI-3**). HC clusters also supported that introversion, extraversion and sexuality are perceptual dimensions, but HC clustering added more fine-grained social context for these dimensions. The features in the cluster Introversion were identified as introvert (PC5), not dominant (PC2), cognitive (PC3) and disengaged (PC4) while the features in the cluster Extraversion & playfulness loaded as extrovert (PC5) but also pleasant (PC1), dominant (PC2), physical (PC3) and playful (PC6). Features in the sexuality cluster loaded as sexual (PC6), pleasant (PC1), empathetic (PC2) and physical (PC3).

While most of the HC clusters followed semantically the main dimensions from PCoA, HC also identified clusters for different communication types (Social engagement, Emotional expression, Gesturing and Physical affection), body movement, physical discomfort, achievement and motivation. These clusters combined information from multiple main dimensions and had semantically distinct meaning not related to any main dimensions suggesting that semantically independent social clusters can be perceived as complex associations from multiple main perceptual dimensions. See interactive 3D representations of the social perceptual space based on PCoA loadings and HC clusters (https://santavis.github.io/taxonomy-of-human-social-perception/Media_social_perceptual_space_3D/social_perceptual_space_comp1-comp3.html).

## Study 3

The purpose of Study 3 was to test whether social perceptual dimensions identified in Study 2 generalize across perceivers and stimuli. As a validation dataset we used a previously collected data where an independent set of participants watched a full Finnish movie in short clips and evaluated the presence of most of the same social features than in the primary dataset. Dimension reduction analyses designed for Study 2 were independently applied for the primary dataset and for the validation dataset. Then the similarity of the identified taxonomies was tested statistically between the datasets.

## Method

As the primary data we used the full rating dataset from Study 1 thus the stimuli, participants and evaluated social features were the same as described in Study 1 methods.

### Evaluated social features

The validation dataset was acquired previously for an independent study and contained most of the social features (76 out of 138) also evaluated in the primary dataset.

### Stimulus

The primary data were compared against perceptual ratings from a slightly shortened version of a Finnish feature film “Käsky” (Aku Louhiniemi, 2008, ∼70min). The movie was spoken in Finnish while the primary dataset with unrelated movie clip dataset contained only spoken English, thus the structural and linguistic variation ensured that we tested the generalizability across distinct data types.

### Participants

The social features in the validation dataset were collected in Finnish from five independent local Finnish subjects not participating in the primary dataset rating. All subjects were female undergraduate psychology students.

### Data collection

The full-length movie in the validation experiment was split into interleaved eight second clips. The participants watched and rated the clips in chronological order. Interleaved data collection resulted in four second temporal accuracy in the validation dataset while each movie clip in the primary dataset was rated only once (average duration of the movie clips: 10.5 sec). Before dimension reduction analyses the ratings were averaged over the participants within a dataset.

### Generalizability of the social perceptual structure between movie stimuli

We tested the generalizability of the PCoA and HC dimension reduction solutions for the organization in social perceptions between the movie datasets. For both analyses the initial dimension reduction was conducted separately for both datasets similarly as described in the section “Low-dimensional model for social perception**”** under Study 2. For PCoA we then identified how many components were statistically significant in both datasets as in the main analysis in Study 2 (null distribution generation with 1 000 0000 permutations) and correlated the feature loadings of the significant components between the datasets. Statistically significant correlations of component specific feature loadings between the datasets would indicate that these components share similar information. For HC analysis the similarity of the cluster structure between the datasets was tested by comparing the consensus matrices as well as the correlation matrices of the datasets with a non-parametric Mantel test with 1 000 000 permutations (Glerean et al., 2016).

## Results

### Generalizability of the social perceptual structure between movie stimuli

To test for the generalizability of the low-dimensional structure for social perception between different movie stimuli and participants, we first conducted PCoA and HC analyses for an independent dataset where a full-length movie was used as stimulus and compared those with the primary dataset. Both PCoA and HC analyses showed statistically significant generalization between the datasets (Figure 7). Permutation tests for the PcoA analyses showed that the movie clip dataset with 76 social features (features common to both datasets) could be described with seven PCs and the full movie dataset with ten PCs (PCs with p < 0.05). Figure 7a shows the statistically significant correlations of the PC feature loadings between the two datasets. Each PC in the primary movie clip dataset showed significant correlation with a corresponding PC in the validation full movie dataset indicating that the corresponding PCs shared similar information. For HC analysis a representative number of clusters was addressed independently for both datasets (**Figure-SI4**). Based on the Mantel test for similarity of two independent matrices the covariance structure between social features was highly similar across the datasets (Correlation matrices: r = 0.68, p < 10^−6^, Consensus matrices: r = 0.54, p < 10^−6^, Figure 7b).

**Figure 7.**
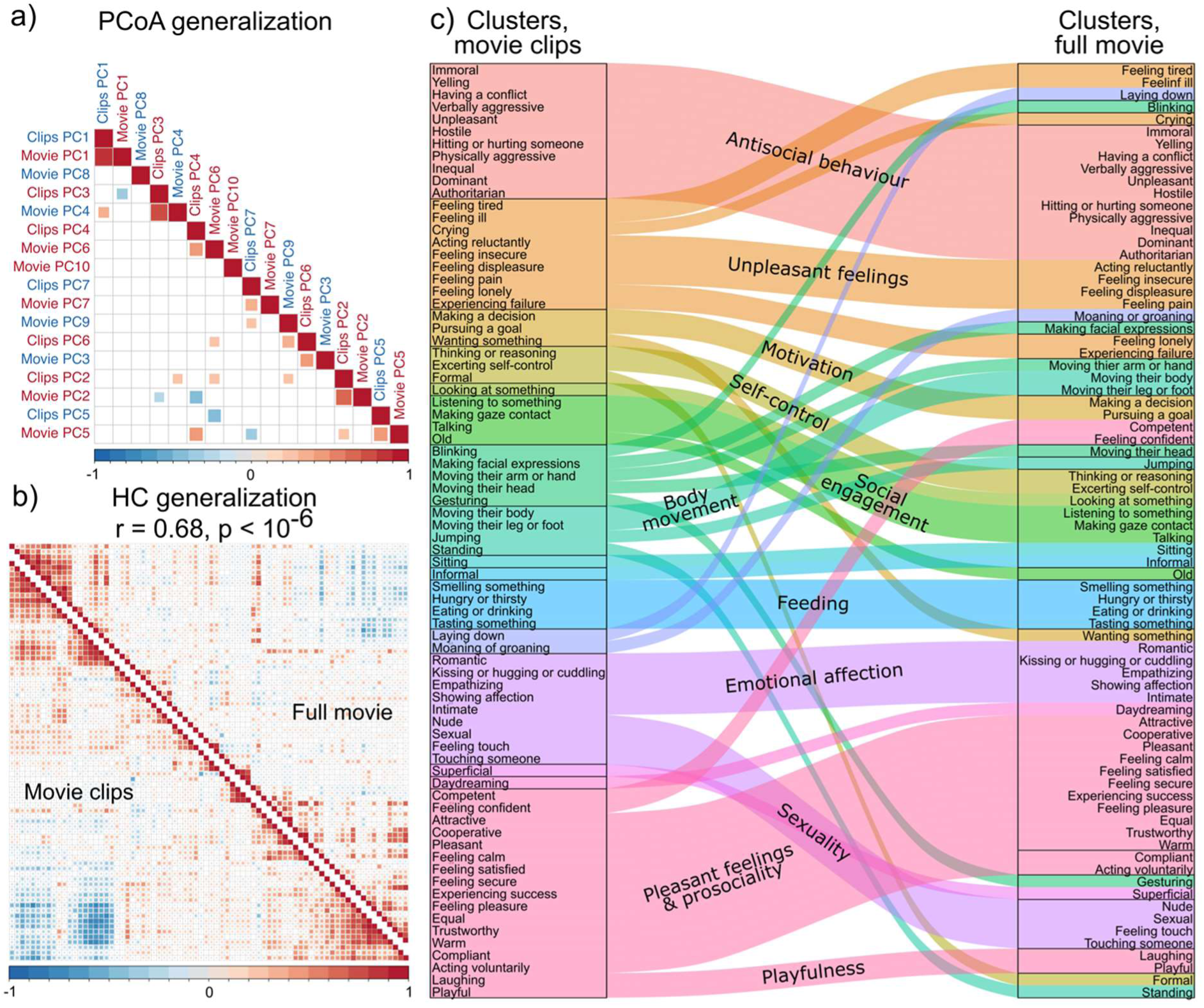
Generalizability of social perception from short movie clips to full movie stimulus. A) Correlations of the PC feature loadings between the independent datasets. Only statistically significant (p < 0.05) correlations are shown. B) Hierarchically ordered correlation matrices from HC analysis. The clustering result showed statistically significant (p < 10^−6^) correlation between independent datasets. C) The alluvial diagram shows the similarities and differences of the clustering results from the independent datasets in detail. The order and grouping of the individual features follow the clustering results obtained independently for both datasets.

## Study 4

Human social perception is inherently dynamic, and it occurs across multiple overlapping temporal scales. However, a large bulk of psychological studies has been conducted using static stimuli not necessarily representative of complex natural world. Hence, the purpose of Study 4 was to investigate how similarly people perceive and evaluate static and dynamic social scenes. To that end, we contrasted the perceptual organization of the movies in the primary dataset against representative static frames extracted from the movies. Hence, Study 4 tests the generalization of social perceptual structures across stimulus types (dynamic videos versus statics frames) which however convey partially similar information. Since movie clips provide more information than static frames, we hypothesized that the agreement of the ratings between participants would be higher when participants watch movie clips compared to watching captured frames from the same movies. We also hypothesized that the proposed taxonomy for social perception would generalize well across stimulus types (Hypothesis 3).

## Method

### Stimulus set

The movie clip stimulus was the same as used in Study 1. A research assistant not familiar with the project was instructed to capture two frames from each movie clip. The only instructions given were that the contents of the captured image should be clearly visible and if the movie clip contains multiple scenes the two images should be from different scenes to ensure variability in the images. A total of 468 images were extracted from the primary movie clip stimulus set.

### Data collection

The data for the image dataset was collected with the same design than the movie clip dataset by replacing the movie clip stimulus with the captured images in the experiment platform Gorilla (https://gorilla.sc/). Each participant now evaluated 78 images instead of the original 39 movie clips. The median experiment duration for the participants was 30 minutes and 28 seconds.

### Participants

The primary movie clip dataset was acquired for Study 1. Participants for the image dataset were recruited through Prolific (https://www.prolific.co/), not allowing the same participants to take part in the experiment that were recruited to the movie clip experiment. Otherwise, the recruitment and data quality screening were identical to those described in the section “Participants” under Study 1. New participants were recruited until 10 reliable ratings were obtained for each social feature and image. Altogether 1109 participants completed the study, and 15 (1,4 %) participants were excluded based on the data quality control. The final dataset included 1094 participants from 56 nationalities with over 654 000 data points. 448 participants were female (41 %) and the median age of the participants was 32 years (range: 18 – 77 years). The reported ethnicities were: White (770, 70.4 %), Black (194, 17.7 %), Mixed (55, 5.0 %), Asian (51, 4.7 %) and Other (20, 1.8 %). See **Table SI-3** for the nationalities of participants.

### Comparing the perceptual ratings between movie and image stimulus

We calculated the intra-class correlation coefficients and bimodality measures (bimodality coefficient & Hartigan’s dip test) for the static frame dataset similarly as in Study 1 (section “Analysing how people perceive social features”). ICCs were then compared between the datasets to investigate whether there are significant differences in how coherently people evaluate movie clips and movie frames. Pearson correlation was used to test how similar gradients from low-agreement features to high-agreement features the datasets produce. Two-sample t-test was used to test the overall difference in ICCs between the two datasets. Bimodality tests were used to assess the generalizability of the categorical versus continuous perception. Pearson correlation of the bimodality statistics and intersection of the significantly bimodal distributions are reported as the measures of agreement in the categorical perception between the datasets.

### Generalizability of the low-dimensional model for social perception between movie and image stimulus

We tested the generalizability of the PCoA and HC dimension reduction solutions between the movie clip and movie frame datasets using the same methods described in Study 3 but now all 136 features were common for both datasets.

## Results

### Similarity of the perceptual ratings between movie and image stimulus

The correlation of the feature specific ICCs between the two dataset was 0.88 (p < 10^−43^) indicating that similar gradient from high to low between-participant agreement was identified in both data. However, the overall between-participant agreement was significantly higher when people watched movies compared to movie frames based on the two-sample t-test (ICC_movies_(mean) = 0.39, ICC_frames_(mean) = 0.34, p(difference) < 10^−10^). Correlation of the feature specific bimodality coefficients between the datasets was high (r = 0.91, p < 10^−54^) while the correlation of the Hartigan’s dip test statistics were moderate (r = 0.4, p < 10^−5^). Bimodality coefficient favoured bimodality of 66 features in the movie clip dataset (see full results in Study 1) and 54 out of the 66 bimodal features were also identified as bimodal in the movie frame dataset. Both, conservative Hartigan’s dip test and BC identified eight social features as bimodal in movie clip dataset and four of these were confirmed in the movie frame dataset. Overall, these measures indicate that the features found to be perceived as bimodal were mostly the same between datasets.

### Generalizability of the structure of social perception between movie and image stimulus

The dimension reduction results were stable when people perceived static movie frames compared to the perception of dynamic movie clips. Each of the eight identified components based on the movie clip data (Study 2) was validated by the movie frame data. Figure 8a shows the statistically significant correlations of the PC feature loadings between the two datasets. Each PC in the primary movie clip dataset showed significant and high correlation with a corresponding PC in the movie frame dataset and only few correlations with other components were observed. Figure 8b shows that after consensus clustering the structure of the correlation matrices were almost identical between the datasets (r = 0.92, p<10^−6^).

**Figure 8.**
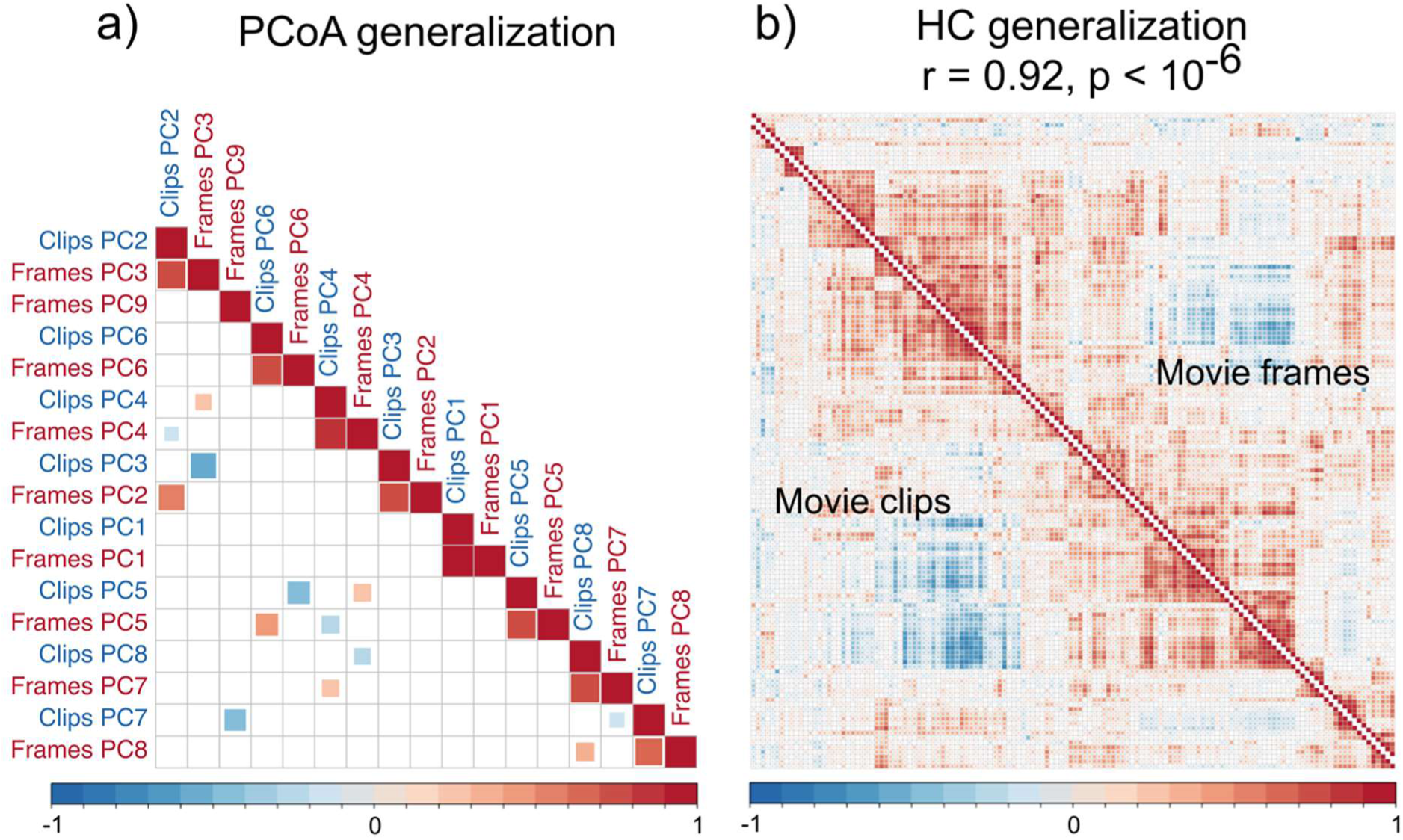
Generalizability of social perception from short movie clips to movie frames from the same stimulus. a) Correlations of the PC feature loadings between the datasets. Only statistically significant (p < 0.05) correlations are shown. B) Hierarchically ordered correlation matrices from HC analysis. The clustering result showed statistically significant (p < 10^−6^) correlation between independent datasets.

## General discussion

Our data present a detailed taxonomy of human social perception based on dynamic perception of social movie scenes. Different social features were perceived with varying consistency between people, and social features with the highest between-subject consistency were perceived categorically (present or absent) while features with low consistency as continuous (intensity). Dimension reduction techniques revealed that, based on the movie stimuli, the social perceptual space can be modelled using eight main dimensions:

1. Pleasant – Unpleasant
2. Empathetic – Dominant
3. Physical – Cognitive
4. Disengaged – Loyal
5. Introvert – Extravert
6. Playful – Sexual
7. Alone – Together
8. Feminine – Masculine.

These descriptive names were defined as the consensus over the responses of authors, researchers, online participants and ChatGPT3.5, and for some component names the consensus of the responses was not as clear as for others (**Table SI-5**). Especially, the “Empathetic” end of the second dimension was often named as “Affection” or “Emotion”, indicating subtle differences in how people viewed this component. Similarly, the component “Disengaged **–** Loyal” was also described with names such as “Passive **–** Proactive” or “Selfish – Selfless” and the “Together” side of component seven was also seen as “Approach” or “Motion”. These subtle differences in the suggested names do not significantly alter the interpretation of the dimensions.

Comparative clustering analysis analyses indicated that the higher-level social clusters can be described as combinations of the fundamental eight main dimensions. The taxonomy showed high level of generalization in multiple levels. The social perceptual structure generalized 1) across independent movie stimuli, 2) from perception of dynamic movies to perception of static frames captured from the same movies and 3) across independent sets of perceivers. Consistent generalization supports the consistency of the established framework for social perception across independent dynamic and static movie stimuli, participants, and two languages. Altogether the results establish how social perception is organized around the potential benefits, harms, and predictability of the social interaction, and how processing of the most critical features is facilitated by categorical rather than continuous perception.

### Is social perception categorical or continuous?

We collected the perceptual ratings on a continuous VAS scale to enable analysis of the rating distribution shapes (Figure 2). This revealed that some social features are perceived categorically either present or absent (Figure 3). Identified features for categorical perception were similar when people perceived movies or static images from the movies, indicating that that the results generalize across stimulus types. Presence of simple interaction signals (e.g. talking, making gaze contact, touching someone, crying, yelling, laughing), feelings (e.g. feeling pain, feeling calm), pair-bonding and features related to sexual behaviour (e.g. nude, sexual, kissing/hugging/cuddling, romantic, flirtatious) and femininity and masculinity were perceived categorically. This does not mean that people cannot perceive or distinguish, for example, different intensities of feeling pain or varying degrees of femininity, but instead, on most occasions the detection of these features is sufficient for guiding subsequent actions in the situation.

Categorical perception is common across the sensory systems (Goldstone, 1994). For example, people perceive colours as discrete despite the fact that they can also perceive continuous changes in the colour spectrum (Pilling et al., 2003). Perception of facial expressions is categorical even though the physical changes of the facial appearance between stereotypical facial expressions are continuous and so forth (Fugate, 2013; Sauter et al., 2011). Our results suggest that perception for social features which carry important social information that may require rapid responses based simply on their absence or presence (such as crying, laughing, feeling pain and features describing sexual behaviour) is initially resolved by categorial perception despite intensity information being present in the scene.

### Consistency of social perception across observers depends on what is perceived

The between-subject consistency of the ratings (measured with ICC) revealed a decreasing gradient from higher agreement in simple actions (e.g. talking, standing, eating), external person characteristics (e.g. nude, feminine), and intense feelings (e.g. feeling pain, feeling pleasure), towards lower agreement in subtle person and interaction characteristics (e.g. punctual, being morally righteous, superficial) when people perceived movies or images (Figure 3). Features with high agreement related to simple observable audio-visual features (e. g. hearing voice and seeing person’s lips moving means that the person is talking). The features with high immediate relevance such as emotional scenes may also capture attention similarly across individuals (Nummenmaa et al., 2006) and are hence perceived with higher agreement than features with low relevance. In line with this, emotionally charged features (e.g. crying, laughing, feeling pain, feeling (dis-)pleasure, sexual) also showed high consistency.

Conversely, the features with lower agreement were indirectly inferred characteristics that require multiple perceptual inputs and holistic processing of the social scene (e.g. evaluating the morality of someone’s actions). It is also possible that the judgment of features with low agreement is largely influenced by perceiver characteristics and top-down signals while sensory information contributes minimally to the bottom-up inference (e.g. perceivers disagree with the semantic meaning of the social feature). Alternatively, perceivers may need to accumulate information across longer timescales to evaluate these features, leading to larger variability particularly in the experimental conditions with short stimulus displays. These explanations are however not mutually exclusive. However, the categorically perceived features were generally perceived with the highest consistency between participants suggesting that the simplicity of the perceived feature drives the perceptual consistency.

It is difficult to comprehensively evaluate our consistency estimates against prior studies, as this kind of large-scale social perceptual studies have not been previously conducted. However, the between-participants consistencies measured with Cronbach’s alpha for evaluation of facial features (α_trustworthiness_ = 0.82, α_dominance_ = 0.88, α_attractive_ = 0.83) align well with those reported in previous face evaluation studies (Carré et al., 2009; Jones et al., 2004; Oosterhof & Todorov, 2008). Our data nevertheless revealed that between-participant agreement was significantly higher when people watched movies than static movie frames from the same movies. These findings suggest that dynamic stimulation model yields more reliable responses compared with static images, most likely because the brain has evolved to react and respond to the dynamic and ever-changing social world rather than static snapshots.

### Taxonomy of human social perception based on dynamic movies

Our results established a detailed model of the organization of social perceptual. The initial set of 138 social features was mostly derived from existing, more narrowly focused models of social cognition, as no previous model has tried to integrate certain subsections of social perception (person perception, action perception, situation perception, trait perception, state perception etc.) into a common framework. To our knowledge, our taxonomy is the first one that is derived from dynamic social perception. Previous models are typically based on perception of static images, or the imagined co-occurrence of different features (lexical studies) that either lack the temporal dynamics of social interaction or the actual perception of social situations altogether.

The primary dimension reduction analysis (PCoA) found statistical evidence for eight orthogonal dimensions as a reduced model capable describing most of the variation in the social perceptual ratings of the movie stimuli (Figure 4). The first three main dimensions describing pleasantness, dominance and cognitive versus physical functions already explained 55 % and eight main dimensions altogether explained 78% of the total variation in the correlation structure between the social features. **Pleasant – unpleasant** dimension explained 32% of the variation and described the overall valence of the social scene. Positively valenced situations promote cooperation, while negative valence would indicate threat or harm promoting avoidance or defensive behaviours. This aligns well with the fact that affective value is one of the primary dimensions in perception in general (Zajonc, 1980). **Empathetic – dominant** dimension explained 13% of the variation and described the social hierarchy between people. Empathic characteristics relate to the intimate and empathetic behaviours towards others while dominant behaviours related to the use of power, agency, and abuse. Dominance is an evolutionary social strategy for maintaining social rank (Maner, 2017) and is considered a basic motivation for humans (Schwartz et al., 2012). Dominance relates to the relative rank in the social hierarchy and relates with social competition. Encoding others’ dominance is critical for avoiding conflicts with higher-ranking dominant individuals but also for forming coalitions against the dominant ones. **Physical – cognitive** dimension characterized social perceptual features from physical actions to cognitive functioning. Impulsivity loaded highly in the physical end of this dimension coupling cognitive and controlled behaviour with impulsive and physical behaviours. Hence, the dimension closely resembles the distinctions of “type 1” and “type 2” or automated versus deliberate processes of social cognition described in the dual-process theories of cognition (Evans & Stanovich, 2013). In situations requiring fast actions, impulsive reactions can be advantageous while complex choices benefit from controlled analytic processing. Therefore, perceiving the cognitive processing types of conspecifics is vital for understanding and predicting the actions of others and thus enables people to tune their own actions accordingly.

These three dimensions already provide a simple yet powerful model for rapid social perception. Moreover, similar dimensions have been identified independently in other areas of social cognition. Already in the 1950s, Osgood’s semantic differential stated that overall meaning of words in general can be summarized in three dimensions, valence, potency and activity (Osgood & Suci, 1955) that resemble to some extent our three first dimensions in social perception. Stereotypic content model and dual perspective model of agency and communion, in turn, have proposed warmth/communion and competence/agency as dimensions to categorize people and groups (Abele & Wojciszke, 2014; Fiske, 2018). Warmth was loaded highly pleasant in the **Pleasant – unpleasant** dimension while agency loaded highly dominant in **Empathetic – dominant** dimension revealing that these two previously established dimensions have a close relation with our first two components. Additionally, valence and dominance have been identified as basic evaluative dimensions of faces (Jones et al., 2021; Morrison et al., 2017; Oosterhof & Todorov, 2008; Sutherland et al., 2013), bodies (Hu et al., 2018; Morrison et al., 2017; Tzschaschel et al., 2022), and even people’s voices (McAleer et al., 2014). General valence is considered a main dimension in many models of emotion and social cognition from the circumplex emotion theory (Russell et al., 1989), to the taxonomies of social situations (Parrigon et al., 2017; Rauthmann et al., 2014). This indicates that similar principles that organize elementary descriptions of words generalize also to descriptions about static (images) and dynamic (videos) features of the social world. Our study is the first to show that these dimensions (along with others, see below) constitute of the elementary evaluative dimensions in dynamic audio-visual stimulus model for social perception and they likely play a major role also in real life social perception.

The data identified five additional significant dimensions. **Disengaged – loyal** dimension described whether people were perceived as engaging actively on social situations (e.g. conscientious, loyal, brave) or not (e.g. lay, superficial, selfish) and **Introvert – extravert** dimension related to social interaction styles. **Disengaged – loyal** closely resembles conscientiousness and **Introversion – Extraversion** extraversion in the Big Five and HEXACO personality theories (Goldberg, 1990; Lee & Ashton, 2004; McCrae & Costa, 1987) indicating that established dimensions in personality traits are also routinely perceived from people in dynamic social situations. Prior studies have indeed shown how conscientiousness and extraversion are primarily inferred from body or whole-person perception (Hu & O’Toole, 2023). Interestingly, our result align well with a previous face perception study that found that inferred extraversion and conscientiousness from faces could not be explained by the perceived valence and dominance of the faces but were instead independent from these basic dimensions (Walker & Vetter, 2016). Accordingly, it is likely that the combined perception of faces and bodies is needed for accurate conscientiousness and extraversion perception and that our dynamic video stimulus captured this whole person perception well.

**Playful – Sexual** dimension described playful / friendly characteristics versus sexuality / mating related features. Perceiving playfulness could help people to evaluate whether to form friendships with others. Humour is a tool to make other people laugh and social laughter enhances social bonding highlighting the importance laughter and playfulness for social bonding and forming non-reproductive alliances (Dunbar, 2012; Manninen et al., 2017; Scott et al., 2014). In contrast, perception of sexual features help to evaluate reproductive qualities of potential mating partners, and the human brain extract these features automatically and rapidly (Hietanen & Nummenmaa, 2011; Putkinen et al., 2023). Playfulness and sexuality were opposite ends of a single dimension, suggesting evaluative distinctions between friends and romantic/sexual partners. **Alone – together** dimension simply described whether people were alone or interacting with others.

**Feminine – masculine** dimension pertained to the (fe)maleness of the individuals in the scenes. This accords with face perception studies showing that femininity-masculinity is a stable perceptual dimension (Little & Hancock, 2002; O’Toole et al., 1998) and traditionally femininity and masculinity have been treated as a single bipolar dimension describing the (fe)maleness as a set of characteristics used to evaluate possible mates (Little et al., 2011; Mitteroecker et al., 2015). Knowing the sex of the interaction partner is obviously relevant for a multitude of reasons ranging from sexual preferences to mate competition and establishment of social alliances, which may be different between the same and opposite sex. The Femininity – Masculinity dimension in our study aligned well with youthful-attractive dimension from the face perception model (Sutherland et al., 2020) and did not couple masculinity with dominance (Oosterhof & Todorov, 2008; Sutherland et al., 2013) nor separated the sex characteristics from youthfulness (Lin et al., 2021).

### How the present findings relate with previous taxonomies of social situations and signals

On theoretical level our findings converge with existing yet more limited taxonomies describing social situations and their encoding from linguistic descriptions. First, the DIAMONDS taxonomy describes social situations using eight dimensions, named Positivity, Negativity, Adversity, Intellect, Duty, Mating, Sociality and Deception (Rauthmann et al., 2014) whereas CAPTION taxonomy (Parrigon et al., 2017) has seven dimensions called Positive Valence, Negative Valence, Adversity, Complexity, Importance, Humour and Typicality. Our model based on direct perception of social scenes in movies accommodates with specific dimensions from both models. First, emotional valence (**pleasant – unpleasant** dimension), which was also the most important dimension in our data, is identified in both models. After this, the CAPTION and DIAMONDS dimensions diverge across the dimensions established in the present study. Dimension Intellect (DIAMONDS) resembles the cognitive end of the **physical – cognitive** dimension and dimensions Duty (DIAMONDS) and Importance (CAPTION) the socially proactive characteristics in our **disengaged – loyal** dimension. The dimension **playful – sexual** relates to dimensions Mating (DIAMONDS) and Humour (CAPTION), whereas Sociality (DIAMONDS) relates to the distinction of non-social and social situations in the dimension **alone – together.** Dimension Adversity is described in both taxonomies, but the interpretation of these dimensions is slightly different. DIAMONDS describes Adversity as situations with conflict, competition and victimization, while CAPTION describes adverse situations as generally depleting. Overall, the adversity resembles the dominant characteristics in our dimension **empathetic – dominant**. No corresponding dimensions were identified for dimensions Deception (DIAMONDS), Complexity (CAPTION) and Typicality (CAPTION). In sum, our model for naturalistic social perception highlights how main components in lexically derived taxonomies for psychological situations can also explain perception of actual dynamic social scenarios, but that they cannot completely cover the complex perceptual and inferential space of rapid social scenes.

Our results also bear resemblance to models describing mental state inference in social settings. The 3d mind model (Thornton & Tamir, 2020) focuses on the dimensions in inferred mental states and could serve as a model for broader social perception. The model organizes mental states in three dimensions: valence, rationality and social impact where the valence and rationality relate closely with the dimensions **pleasant – unpleasant** and **physical – cognitive** in our data. Social impact is described as a dimension from highly arousing and social states, such as lust and dominance at the other end to low arousal and non-social states, such as drowsiness and fatigue (Thornton & Tamir, 2020). Our dimension **empathetic – dominant** includes dominant features on the other end and some features related to fatigue on the other, indicating correspondence between the social impact dimension in the 3d mind model. However, our data indicates that dimension **empathetic – dominant** does not relate to arousal, but rather to a distinction between dominant and “cold” characteristics versus empathic and intimate characteristic. “Emotional arousal” was associated with the empathic end of the dimension, instead of the dominant end thereby diverging from the 3d mind model. Additionally, **Alone – together** dimension in our model distinguishes social and non-social situations. Hence, our data suggests that the 3d mind model of mental states may not fully generalize to naturalistic social perception. Indeed, the mental state and trait inferences from naturalistic stimuli are found to be not independent from each other (Lin & Thornton, 2023) further empathizing that, at least, mental state and trait inferences should be studied together when building taxonomy for social perception.

Finally, our model diverges from some existing data-driven taxonomies. The ACT-FAST taxonomy for action understanding describes complex human actions in six dimensions: Abstraction, Creation, Tradition, Food, Animacy, Spiritualism (Thornton & Tamir, 2022). We found a cluster for feeding and eating related features converging with the Food dimension in the ACT-FAST taxonomy and the feeding related features were prominent in **disengaged – Loyal** dimension but otherwise it is difficult to establish similarities with ACT-FAST. This is most likely because our model intends to describe immediate social perceptual dimensions that are used to make fast and broad inferences of the social situations whereas the ACT-FAST taxonomy categorizes complex and detailed actions that are not necessarily related to social interaction. Similarly, taxonomies for human goals (Wilkowski et al., 2020) and for basic individual values (Schwartz et al., 2012) are difficult to clearly connect with the current taxonomy. Hence, it is probable that other peoples’ goals and values are not perceived independently of other important social information.

In sum, the proposed model for rapid social perception agrees with some of the earlier proposed primary dimensions of social perception, most notably the importance of affect and distinction between empathy versus dominance and physical versus cognitive features. A subset of the dimensions established in our model have been previously proposed in circumplex emotion theory, Osgood’s semantic differential, warmth/communion and competence/agency models, 3d mind model, psychological situation models, face perception models and also in personality trait inference. However, here the dimensions are integrated into a unified model where the relations between the dimensions are characterised, and the additional dimensions (disengaged – loyal, introvert – extravert, playful – sexual, alone – together, feminine – masculine) result in a parsimonious model where eight main dimensions can be used to characterise the majority of the variation in the social-perceptual evaluations.

### Concordance analysis of PCoA main dimensions and HC clusters

PCoA revealed that eight basic dimensions are sufficient for low-dimensional modelling of the social perceptual space. Consensus clustering analysis showed high concordance with the PCoA organization for social perception (Figure 5 **& 6**). The analysis revealed clusters that closely resemble the main dimensions in PCoA supporting that people perceive valence, dominance, cognitive functioning, sexuality, playfulness, introversion-extraversion, and femininity-masculinity from life-like social scenes.

Additionally, HC analysis revealed fine-tuned clusters that combined information from multiple main dimensions (**Figure SI-3**). For example, features in the cluster Unpleasant feelings could be explained with a combination of dimensions related to unpleasantness and empathy in the PCoA analysis. Conversely, features in the cluster Antisocial behaviour could be explained with a combination of dimensions related to unpleasantness and dominance. Similar fine-tuning for pleasant features were identified based on their dominance structures: Pleasant feelings and prosociality (solely pleasant), Emotional affection (pleasant + no competition) and Extraversion & playfulness (pleasant + competition). Thus, both pleasant and unpleasant features formed clusters with distinct dominance structures highlighting the joint importance of valence and dominance.

HC also identified clusters with no clear semantic link with the main dimensions extracted in the PCoA. Instead, the features in these clusters loaded significantly on multiple PCoA dimensions and introduced semantically distinct social information not identifiable from the main dimensions alone. For example, HC grouped communication types into distinct clusters (Social engagement and Emotional expression, Gesturing, and Physical affection) and identified clusters Achievement and Motivation that formed from characteristics often attributed to driven and high-achieving individuals. Overall, the concordance analysis indicated that while social perception can be modelled with fundamental basic dimensions from PCoA, fine-grained social clusters arising from combined information from multiple basic dimensions can be identified with HC.

We used consensus hierarchical clustering approach to find stable social clusters within our data. This does not mean that these are the only meaningful social cluster that can emerge from the basic perceptual dimensions. The purpose of HC was to validate the usefulness of the main perceptual dimensions as a model for immediate social perception and to suggest that people may only need to perceive the main perceptual information of the situation and be still able to make higher-level inferences of the situations with the combined information of the main dimensions.

### Eight basic dimensions of social perception

Based on the four studies and the conceptualization of social perception offered by the dynamic interactive theory of person construal (Freeman & Ambady, 2011) we propose a model where immediate perception of social scenes is based on eight basic dimensions (Figure 9). Low-level sensory information is rapidly extracted and processed. Then, the integrated low-level information is evaluated along eight primary dimensions for guiding immediate social reactions. This rapid evaluation of social situations in a limited set of main dimensions would allow humans to achieve accurate enough interpretation of the situation in limited amount of time preserving the possibility to react swiftly. Subsequently, the basic social information is used for establishing more fine-grained model of the social scenario, which can then be used for adjusting behavioural repertoires during the unfolding social event. This process is dynamic and influenced by situational and person-level factors and constantly updated information is used for guiding subsequent perception and action.

**Figure 9.**
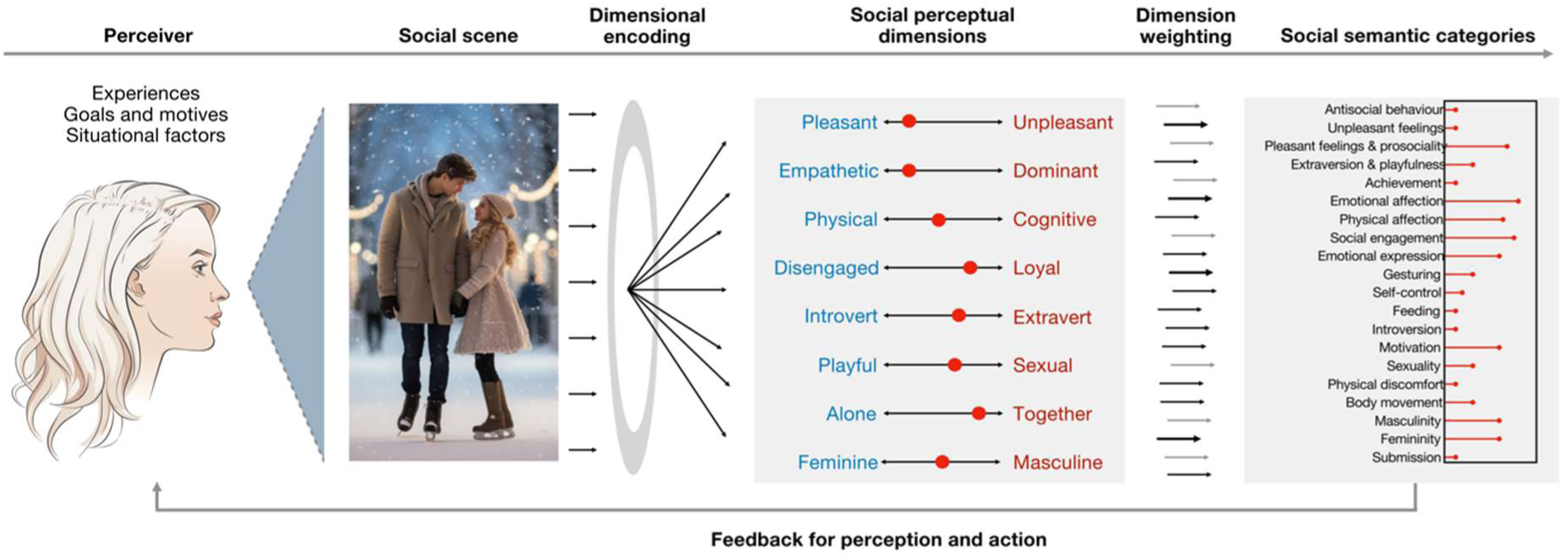
Conceptual summary of social perception based on Studies 1-4. Eight basic social perceptual dimensions are encoded from the low-level information of the social scene. These dimensions can be subsequently integrated for establishing fine-tuned and behaviourally relevant social semantic categories, that may guide action and subsequent perception of the evolving social scene.

We stress that the model describes the population level principles underlying social perception and is based on the primary stimulus set short Hollywood movie scenes. It does not focus on individual differences in perception. In reality, the perceiver and the environment are in a dynamic interaction and perception and interpretation of specific features and dimensions is dependent on perceiver characteristics, such as their experiences, values, goals and emotions (Hehman et al., 2017; Judd et al., 2019). Thus, individuals may perceive social situations differently and the within subject dimensionality in the perception can differ from our model for the population level average social perception. Accordingly, one interesting avenue for further research would thus be to compare the social perceptual dimensionality between healthy participants and patients with neuropsychiatric conditions.

### Advantages and disadvantages of using movies as stimulus model for social perception

We used movies as a stimulus model for life-like social perception. Movies have the advantage of being highly natural, dynamic and high-dimensional stimuli while still offering high degree of experimental control to study social perception (Adolphs et al., 2016), emotions (Saarimäki, 2021), psychiatric illnesses (Eickhoff et al., 2020) or neurodevelopment (Vanderwal et al., 2019). There is also evidence within social neuroscience that the perceptual results from extensively used static images do not generalize fully to those derived from dynamic stimulus. For example, face selective areas in superior temporal sulcus (STS) respond more strongly to dynamic (artificially dynamic or movie stimulus) than static faces (Kessler et al., 2011; Pitcher et al., 2011) and numerous studies, in addition to our previous studies (Lahnakoski et al., 2012; Santavirta et al., 2023), have mapped social function for STS (Lee Masson & Isik, 2021; Nummenmaa & Calder, 2009; Saitovitch et al., 2019). Together these findings have revealed a potential visual pathway that encodes dynamic aspects of social perception (Pitcher & Ungerleider, 2021) suggesting that dynamic stimuli may reveal life-like social processing more reliably than static images (or written vignettes or lexical comparison of words that do not require external perception at all). Consequently neural responses to naturalistic movies are more reproducible and reliable than those to static images or artificial stimuli (Eickhoff et al., 2020; Hasson et al., 2010; Nastase et al., 2020; Sonkusare et al., 2019) and social neuroscience have been increasingly using movies as stimuli to study socioemotional processes in the human brain (Hasson et al., 2004; Lahnakoski et al., 2012; Nummenmaa et al., 2023; Saarimaki et al., 2016; Santavirta et al., 2023; Wagner et al., 2016). Movies are engaging, and a wide range of social situations can be shown in limited amount of time. Movies can also safely depict alarming social situations that are highly relevant from survival perspective (e.g. violence) making them a powerful stimulus model for social perception in the laboratory setting.

Hollywood movies, as basically all previously used stimuli for studying social perception, have also disadvantages. Movies often present stereotypical or “amplified” versions of social interactions that are presented with high frequency, which may lower the generalizability of our findings to real life social events. Such amplification is, however, not only a disadvantage since it allows studying the whole intensity spectrum of the studied phenomenon, and the fast frequency of the socioemotional episodes is a major advantage for generating efficient stimulation models for the high-dimensional stimuli in psychological experiments. To limit the most obvious biases, the presently used movie clips were chosen to depict scenes possible in real life e.g. exaggerated emotions or actions were not included. Movie clips typically present intense scenes, but the movie clip set contained both mundane social events and intensive social scenes to cover social perception in broad intensity spectrum (**Table SI-2**).

Hollywood movies may depict sex stereotypes (Kumar et al., 2022). Sex bias and other possible biases regarding the variation in movie characters features (such as ethnicity, age distribution, attractiveness, body composition …) are widely discussed in the public and they can affect peoples’ perceptions, but there is actually little empirical work on whether these potential biases influence perception differently in Hollywood movies versus real life social scenarios that also involve stereotyping of people. Many of the movie clips in the primary dataset contained actors that are famous around the world (e.g. Leonardo DiCaprio), while a large proportion of scenes did not contain famous actors. We did not measure the familiarity of the actors to the participants or whether the participants had seen the movies before. Hence our results are influenced by the population level familiarity of the actors / movies and prior impressions of the actors and first impressions to the particular short movie scenes cannot be untangled. In real life, people engage in social situations that are familiar to them to some extend (e.g. I know the people in the situation or I have been in a similar situation before) where it is also not possible to differentiate naïve first impressions from prior knowledge. Finally, we used short (∼10sec) movie clips for studying immediate social perception in dynamic scenes. However, social perceptual information may unfold in longer timescales and these results cannot capture those slow perceptual processes.

To quantify the possible biases in out stimulus set we quantified the age, sex and inferred ethnicity of the characters depicted in the movie clips. The clips depicted people from both sexes, from different age groups and multiple ethnicities (118 clips contained females, 197 males, 77 children / adolescents, 192 adults, 10 older adults, 213 Caucasians, 44 African Americans, 10 persons from East Asia and 4 persons from the Middle East). We acknowledge that older people, women, and people of colour are less well represented in this set of stimuli than their distribution in the population in planet earth. It is known that people categorize others quickly by their ethnicity (Kubota & Ito, 2007), and the ethnicity of the target or the perceiver can influence emotion recognition accuracy (Elfenbein & Ambady, 2002; Halberstadt et al., 2022; Mende-Siedlecki et al., 2021). Hence, the ethnicity of the movie characters may have influence on the findings. However, we did not find evidence that the demographic factors (age, sex and ethnicity) of the participants influenced their perceptual rating of the stimulus movies (Figure 10). As a complementary analysis we quantified whether the similarity of the social perceptual ratings would associate with demographic factors of the participants (see section “Associating perceiver’s age, sex, and ethnicity with the social perceptual ratings” in supplementary materials for analysis details). We did not observe major differences in the correlations between individual participant’s ratings compared to others by age, sex, or ethnicity. The correlation between age and the participant’s rating correlation with others was 0.02. The mean rating correlation with the same and opposite sex perceivers was 0.58 and 0.57, respectively. The mean rating correlations by ethnic groups were also highly similar (Asian: 0.60, Black: 0.57, Mixed: 0.59, White: 0.63, Other: 0.65), confirming that at least these features contribute minimally to perception of the social scenes in the movies.

**Figure 10.**
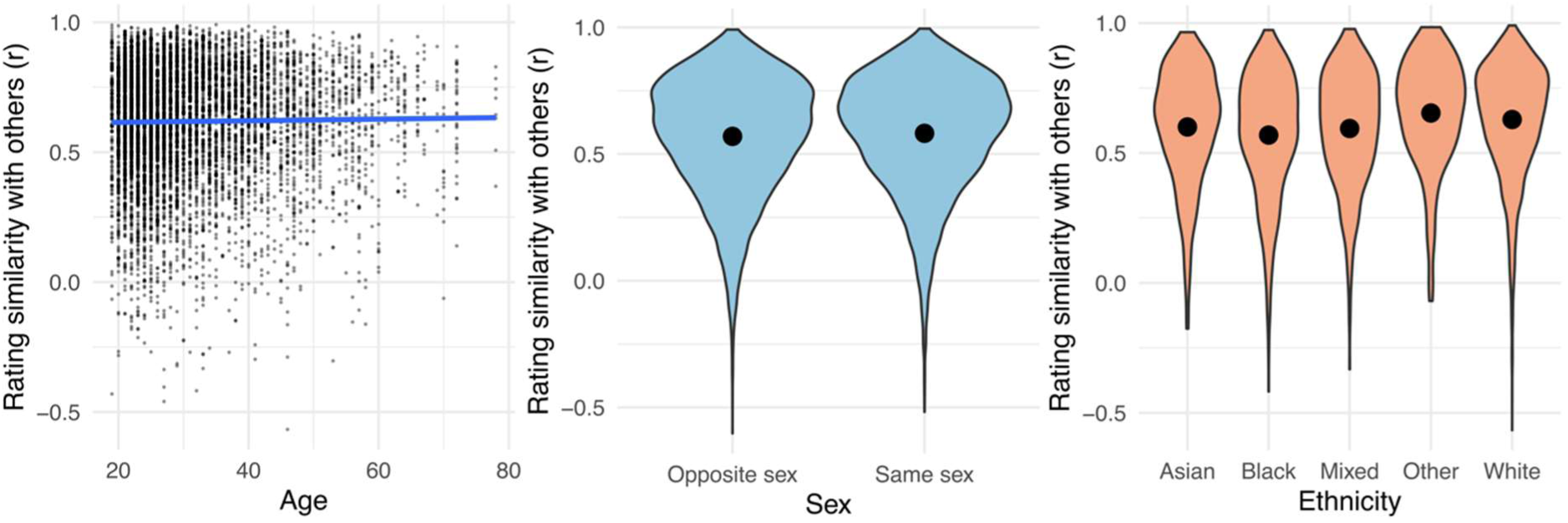
Individual participants’ ratings correlation with the mean of other participants’ ratings based on the participants’ age, sex and ethnicity. Blue line in the scatterplot visualizes the linear association between rating similarity and age. The black points in the violin plots mark the group means.

Considering the evident advantages of movies, the breakdown of the demographics of the movie characters, the lack of major influence of perceivers’ demographics to the perceptual ratings, and the robust generalizability of the derived social perceptual model to independent full movie stimulus, we conclude that the used movie clip dataset was a capable stimulus model for dynamic social perception. As with all experimental designs, this stimulus model comes short from real social life of humans as discussed above. As a form of art of media, the

### Feature selection in data-driven studies of social perception

The current model for social perception is based on data-driven analyses of high-dimensional dataset including 138 pre-selected social features. The identified dimensionality is dependent on the social features selected for evaluation and the feature set also affects the estimations of the relative importance’s (variance explained) of the PCoA dimensions. The included social features may not encompass all possible social scenarios and therefore our model does not rule out the existence of other social perceptual dimensions. The feature set was compiled based on existing theories of social perception as described in the Method of Study 1 with the goal to be comprehensive enough to capture important elements of social perception without being too exhaustive for reasonable data collection. We acknowledge that the feature selection required researcher-based decisions which may influence the resulting findings, similarly as all studies in experimental psychology where dimensional ratings are acquired without open labels (e.g. facial expression recognition, attitude evaluations or responses to personality test items). We however stress that the results also generalized to another movie stimulus as well as static images, indicating that the dimensionality with this feature set replicates across different stimuli. Some studies have freed the researcher from active decisions in feature selection by letting participants spontaneously define the scales and dimensions (Koch et al., 2016; Nicolas et al., 2022; Osgood & Suci, 1955). This decision shifts the feature selection from researchers to the participants which could however result in a bias towards conscious reasoning and overlook unconscious processing of important social features which may be better captured when building the feature set based on the existing theories.

### Future directions

The current research opens numerous exciting avenues for future research. First, sex, ethnicity, familiarity or attractiveness of the actors and other biases in Hollywood films may have influenced the current results and it would be highly interesting in future to test broader groups of dynamic stimuli and participants or to directly test the influence of different stereotypes on dynamic impressions. Results based on Hollywood movies could be tested with stimuli containing Bollywood / Nollywood cinema, TV talk shows, documentaries, or recordings of live theatre, to investigate the social perception structures with other types of dynamic media. Ultimately, future research should push for replicating our results with real-life recordings of human social interactions captured with action cameras by volunteers from their everyday life. Future studies should also move away from the “spectator” perspective by allowing people to engage in social situations instead of passive observing. However, with this kind of real-life stimuli it would be unethical to investigate many of the perceptual categories (e.g. violence, sex) tested in the current study. Second, the dimensionality established with the current theory-driven choice of rated social features could be compared to a high-dimensional dataset where social features would be pooled from a set of naïve participants thereby leaving the choice of the rated features to study subjects. One intriguing line of research would be to test the current abilities of large-language-models (LLMs) for i) generating dynamic video stimuli with predefined features, ii) pooling large and diverse sets of social features for humans to evaluate, or iii) even using LLM as a perceiving participant for social feature evaluation. This type of research would obviously be limited by the possible biases of the training data of the LLM, but LLMs have the advantage of massive training datasets going beyond what could ever be achieved using human observers. Finally, the present results describe only moderately fast social perception in the timescale of ∼ 10 seconds. It would be highly interesting to study how the perceptual structure of social features and the evaluation consistency between participants would evolve with shorter (milliseconds) or longer (minutes) time scales. Again, such work could be complemented by the LLMs by limiting the temporal information given to the model.

## Conclusion

Our results establish a representation of how rapidly people perceive social situations. Social signals are perceived categorically (either present or absent) or continuously (how much). Simple social features and the features with immediate social relevance are perceived with the highest consistency between people indicating that perceptual simplicity and immediate social relevance drive the consistency of perception between people in naturalistic observation of social scenes. Social perception in movie stimulus model organizes into eight main perceptual dimensions that can describe the rudimentary social information of the scenes. Emotional valence, dominance versus empathy, and cognitive functioning versus physical behaviour are the most fundamental perceptual dimensions and they explained over half of the variance in the whole data. Distinct social clusters can be represented based on the join information from the main perceptual dimensions suggesting that basic social information is filtered from social cues and that basic information from multiple dimensions can be integrated to identify more detailed social information. The established model for social perception generalized across stimuli and participants validating the findings. Based on the results, we propose **eight basic dimensions of social perception** model for rapid social perception where social situations are perceived along eight orthogonal perceptual dimensions (most importantly emotional valence, empathy versus dominance, and cognitive versus physical behaviour) and whose combinations can be used for describing detailed qualities of social situations.

## Ethics statement

University Turku Ethics Committee Humanities and Social Sciences Division waived the study from ethical review due to minimal impact on human subjects. All participants gave an informed consent and were compensated for their participation. We report how we determined our sample size, all data exclusions, all manipulations, and all measures in the study.

## Data and code availability

The anonymised perceptual rating data and analysis scripts are available in the project’s Git (https://github.com/santavis/taxonomy-of-human-social-perception)(Santavirta, 2024). The stimulus movie clips can be made available for researchers upon request, but copyrights preclude public redistribution of the stimulus set. Short descriptions of each movie clip can be found in the supplementary materials (**Table SI-2**).

## Author contributions

SS designed the experiment, collected data, developed the analysis methods, analysed the data and wrote and reviewed the manuscript.

TM designed the experiment and wrote and reviewed the manuscript. AE collected data and wrote and reviewed the manuscript.

LN conceptualized the study design, acquired funding, supervised the project, developed analysis methods, and wrote and reviewed the manuscript.

## Acknowledgements

The study was supported by Academy of Finland grant (#332225) to LN, Turku University Foundation and Alfred Kordelin Foundation grants to SS, Finnish Governmental Research Funding for Turku University Hospital and for the Western Finland collaborative area to SS and TM and Päivikki and Sakari Sohlberg Foundation, Finnish Brain Foundation grant and Finnish Cultural Foundation grants to TM. We thank Tuomas Knuuti for his help with data collection.

