## Supplementary materials for "A taxonomy for human social perception: Data-driven modelling with cinematic stimuli"

### Supplementary methods

#### Experiment subsets

Based on factors in situation taxonomies we hypothesized that emotional valence would be a main dimension of social perception (Oosterhof & Todorov, 2008; Parrigon et al., 2017; Rauthmann et al., 2014; Santavirta et al., 2023). To ensure that each participant would evaluate a subset of all movie clips with a wide range of social interaction and balanced emotional content, the movie clips were divided into three broad sets based on their emotional valence (negative / neutral / positive). The clips were then divided into six subsets of 39 clips where each contained an equal number of clips from negative, neutral, and positive emotion categories. Social features were then divided into 18 sets (6-8 features within a set) so that each set contained only features from a single category (e.g. person's traits, physical characteristics…). Upon recruitment each participant was randomized to evaluate one set of social features (6-8 features) from one set of movie clips (39 clips).

#### Data quality control

We excluded subjects based on the following data quality criteria: 1) A participant had not moved the slider at all in at least 50 % of the video clips. 2) A participant had not moved the slider below 50 (average value) in any video clip. 3) A participant had not moved the slider below 50 in at least 5 consecutive videos in the beginning of the survey (indicating misunderstanding of the instructions). 4) A participant only gave minimum (0) or maximum (100) ratings for each video clip and every social feature. Criteria 1-3 were evaluated separately for each rated social feature and a participant was excluded if the data quality was considered poor in even one of the rated social features.

#### Sample size estimation

To estimate how many participants were required for the primary movie clip dataset, we studied how sample size affects the stability of intra-class correlation coefficient (ICC) estimates in an unrelated dataset where emotion ratings were obtained using a similar setup. This sample of emotion ratings included 27-28 independent ratings for 33 emotion features. For each possible subsample of the ratings (from 2 to 26) we sampled 1000 subsets (or all possible sets, if possible sets < 1000) from the original data and estimated how much the ICC varies between the subsets for each sample size. The variation in ICC between subsamples was calculated as the width of 95% quantile interval ($Q_{97.5\%}$ - $Q_{2.5\%}$) of the ICCs. The increase in the stability of ICC estimates (decrease in variation) was modest when the sample size increased over 10 ratings and therefore sample size of 10 ratings for each social feature and movie clip was considered sufficient. **Figure SI-5** shows the stability of ICC estimates in the emotion dataset as well as the within sample stability of ICC estimates in the collected social dataset.

Associating perceiver's age, sex, and ethnicity with the social perceptual ratings

Our model for social perception was derived from the full dataset (primary movie clip data) including participants with wide age range, from both sexes, and from different ethnicities. Age, sex and ethnicity of the participants were extracted from standard demographic report of the online data collection platform Prolific. To estimate whether perceiver's age, sex or ethnicity influenced how they perceived the movie clip stimuli we analyzed the correlations of the ratings between subjects. To estimate the association of age and ethnicity with social perceptual ratings we calculated the correlation of every single participant's ratings with the mean ratings of all other participants who rated the same social feature from the same movie clips. To estimate the association of sex and social perceptual ratings we calculated the correlation between every single participant's ratings with the mean ratings of all other participant with the same and opposite sex separately. If people with different demographic factors would perceive the movie clips differently, we would expect to observe clear association between the individual rating correlations with others based on the participants' age, sex or ethnicity. **Figure 10** in the main text visualizes how age, sex or ethnicity influenced the participants’ perceptual ratings of the movie clips.

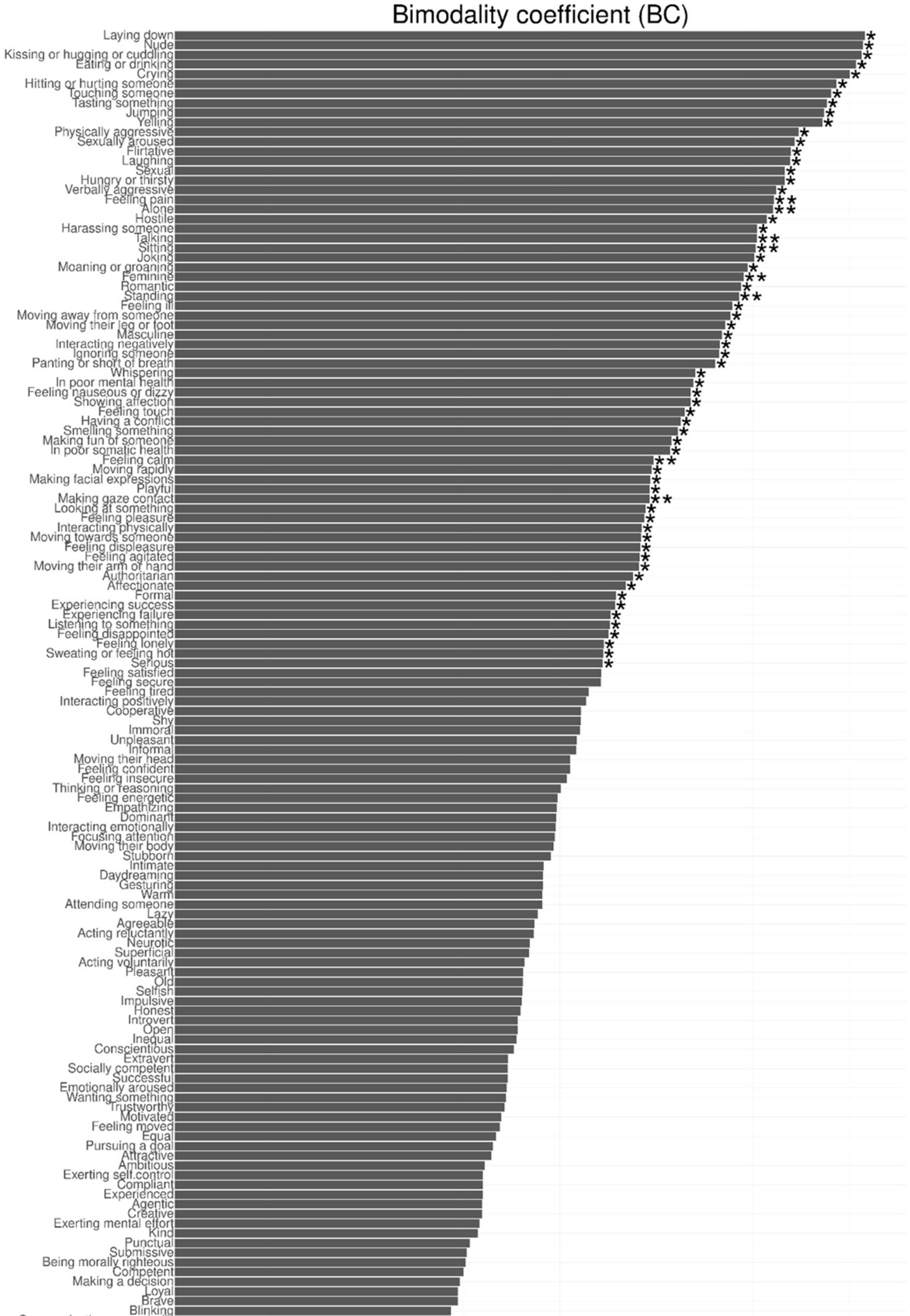


**Figure SI-1** Bimodality analysis results. Bimodality coefficient (BC) and Hartigan’s dip test p-value were used as metrics to evaluate whether on average the people rate social features as unimodal or bimodal. Feature barplots visualize BC of the features. ** marks that the dip test rejects unimodality ($p<0.05$) and that bimodality coefficient favors bimodality ($BC > 0.555$). * marks that only bimodality coefficient favors bimodality ($BC > 0.555$).

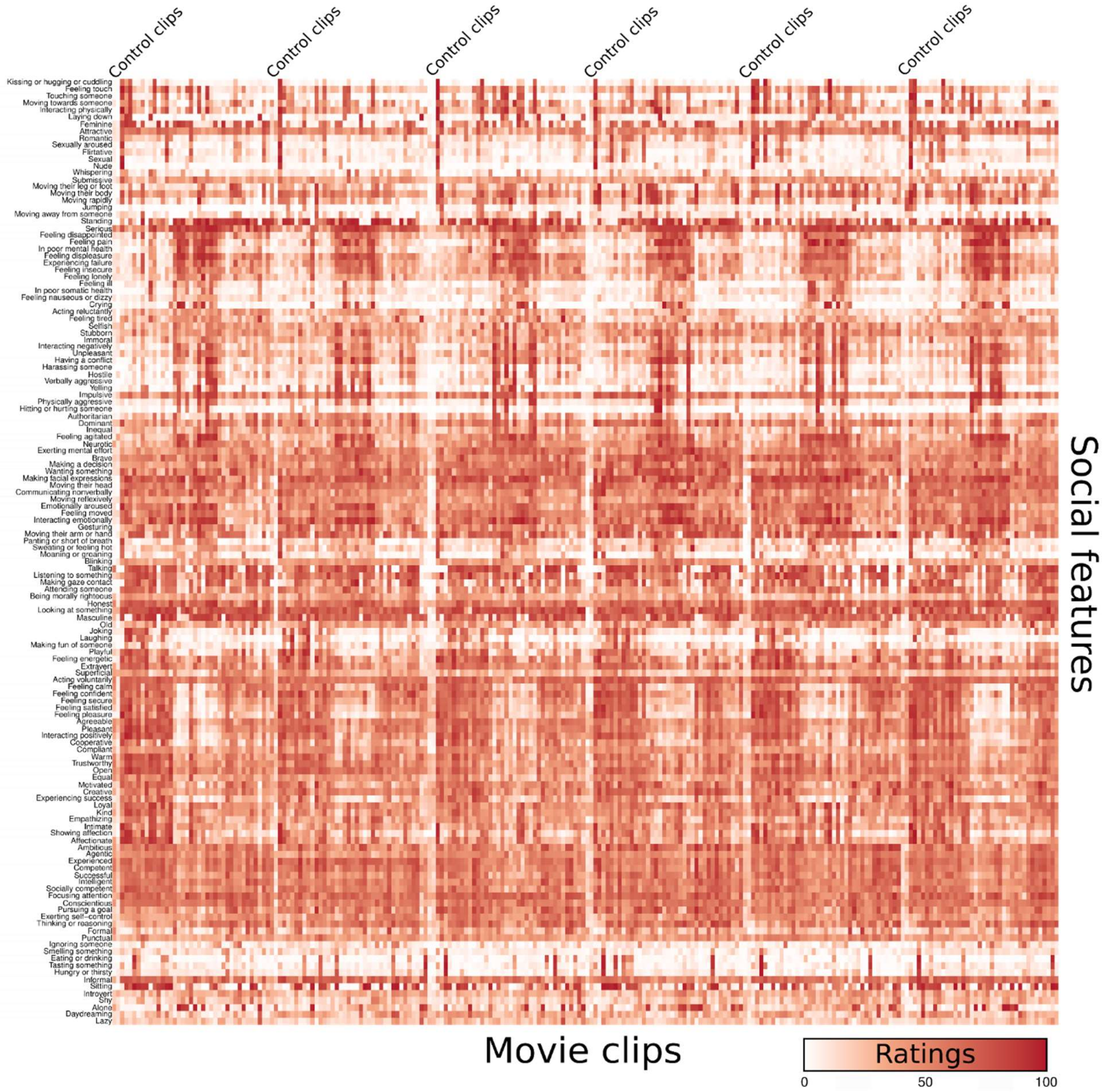


**Figure SI-2** Heatmap of social feature ratings to show the variability in the rating intensities between movie clips. Each cell visualizes the population average rating intensity of the given social feature.

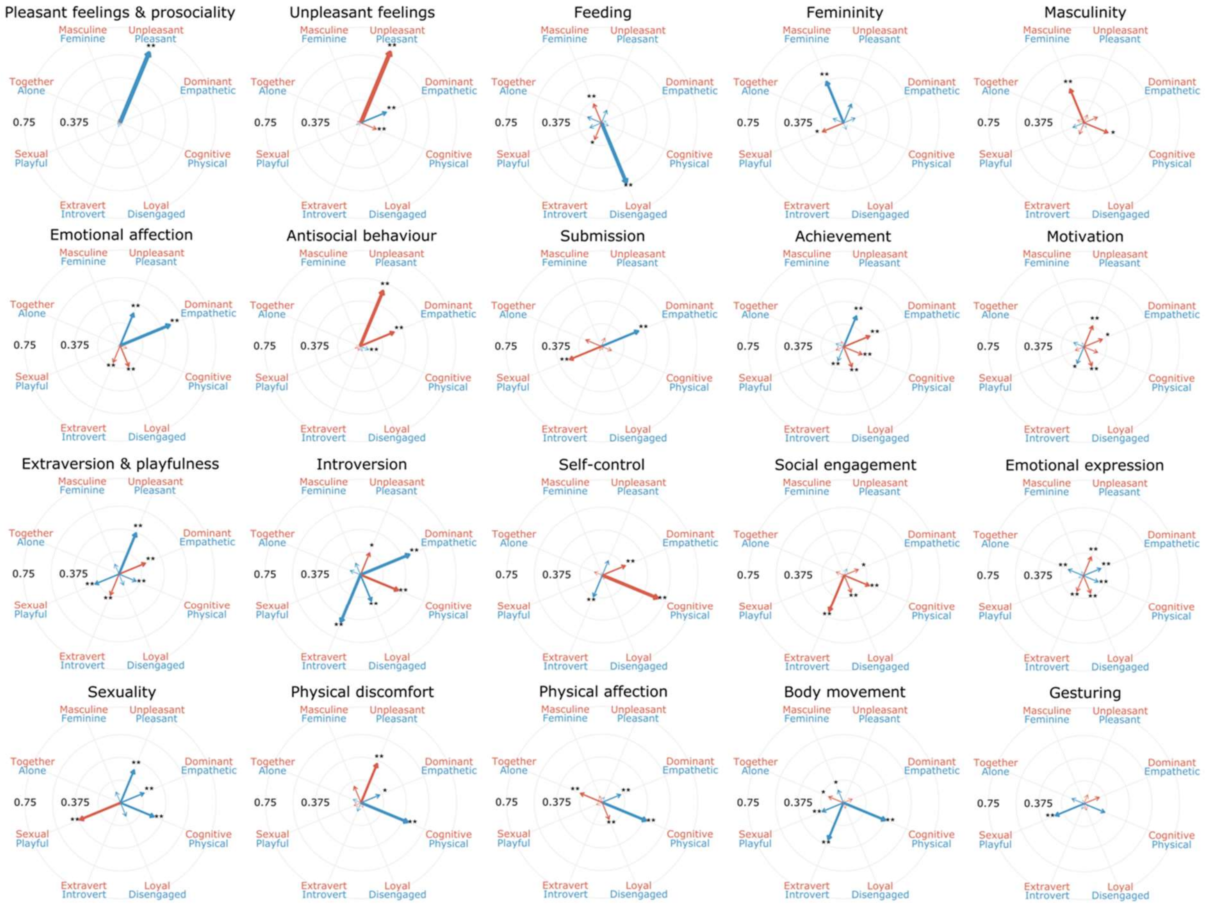


**Figure SI-3** The relationship between hierarchical clusters and principal components from the principal coordinate analysis. The polar plots show how social features on average loaded for PCs in hierarchical clusters. The length of each arrow visualizes the absolute loading of the cluster for a given PC while the color of the arrow marks the direction of the loading. ** marks strong statistical significance of average loadings ($p < 0.05$, Bonferroni corrected for the total number of clusters) while * marks a more lenient statistical thresholding ($p < 0.01$, uncorrected).

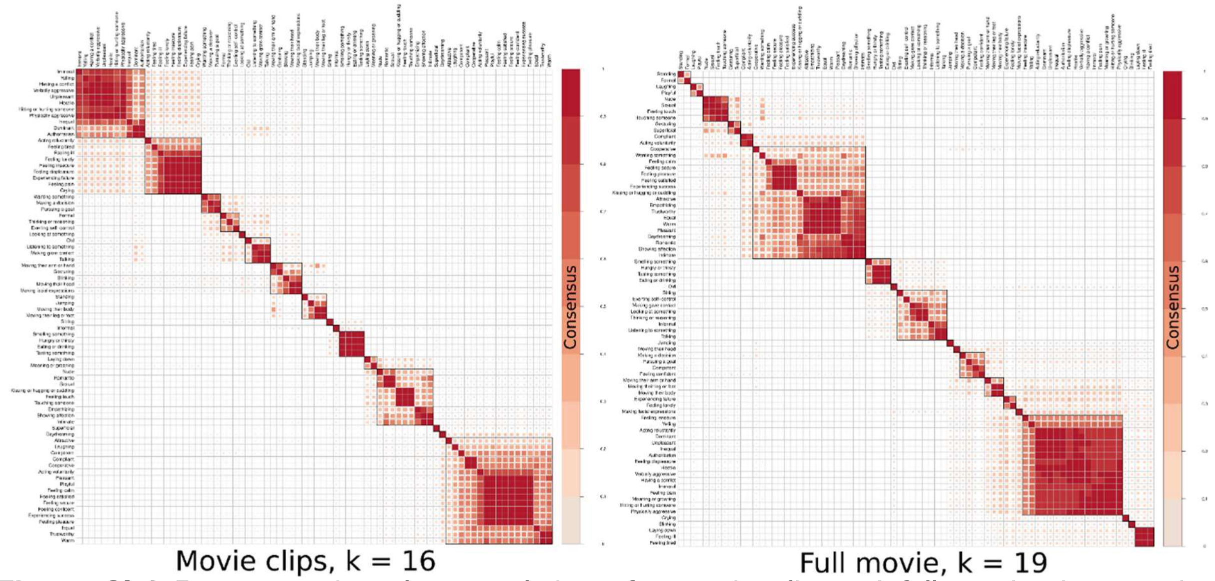


**Figure SI-4** Representative cluster solutions for movie clip and full movie datasets in generalizability analysis. Representative number of clusters were selected for both datasets independently based on their hierarchically ordered consensus matrices.

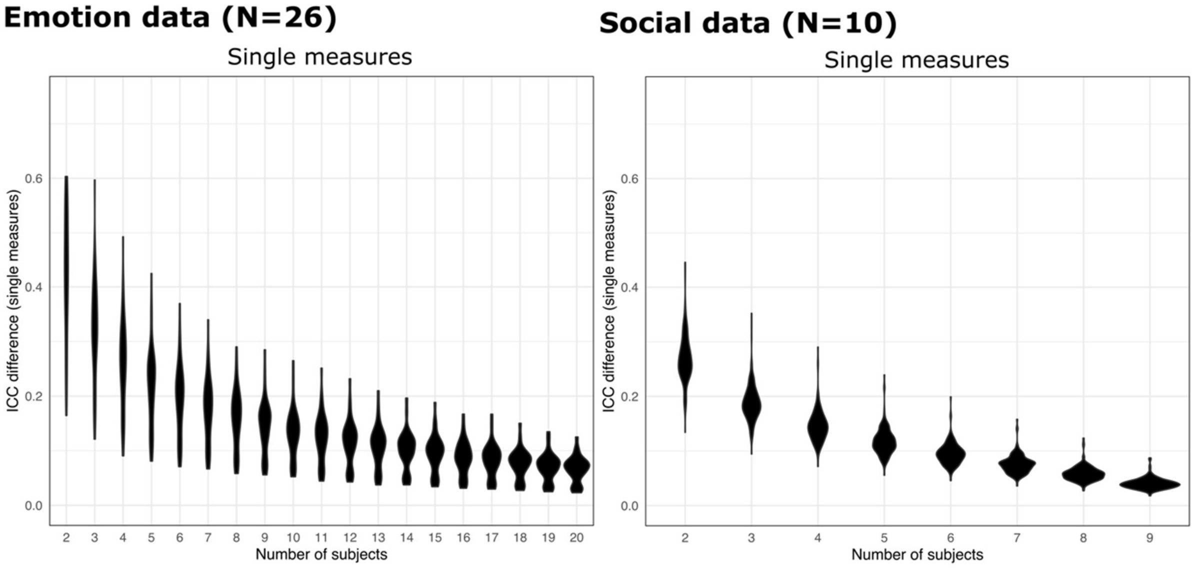


**Figure SI-5** Permuted stability of ICC estimates of socioemotional ratings as a function of sample size. Top row shows the stability of ICC estimates in a previously collected dataset of perceived emotion features in similar study design with a sample size of 27 ratings for each emotion feature and video clip. Single measures ICC reliability is plotted on the left while the right panel shows the average measures ICC reliability. The violin densities reflect how much the ICC estimates vary in the permuted subsamples of all rated emotion features. The bottom row shows the results of a similar analysis from the collected social dataset with 136 different social features.

**Table SI-1** Rated individual social features and their links with the established psychological taxonomies. See *TblSI1_featurestheory.xlsx*.

**Table SI-2** Short descriptions of the movie clips used as primary stimuli. See *TblSI2_clip_descriptions.xlsx*.

**Table SI-3** The nationalities of the participants in movie clip and movie frame datasets. See *TblSI3_nationalities.xlsx*.

**Table SI-4** All outcome results for the individual social features in the movie clip dataset. *TblSI4_full_result_table.csv.*

**Table SI-5** Suggested PCoA component names by authors, researchers, online participants and ChatGPT3.5. *Tbl5_component_naming.xlsx*

**Table SI-6** Suggested HC cluster names by authors, researchers, online participants and ChatGPT3.5. *Tbl6_naming_naming.xlsx*
